# Structure of PSI-LHCI from *Cyanidium caldarium* provides evolutionary insights into conservation and diversity of red-lineage LHCs

**DOI:** 10.1101/2023.10.25.563911

**Authors:** Koji Kato, Tasuku Hamaguchi, Minoru Kumazawa, Yoshiki Nakajima, Kentaro Ifuku, Shunsuke Hirooka, Yuu Hirose, Shin-ya Miyagishima, Takehiro Suzuki, Keisuke Kawakami, Naoshi Dohmae, Koji Yonekura, Jian-Ren Shen, Ryo Nagao

## Abstract

Light-harvesting complexes (LHCs) are diversified among photosynthetic organisms, and their structural variety in photosystem I-LHC (PSI-LHCI) supercomplexes has been shown. However, structural and evolutionary correlations of red-lineage LHCs are unknown. Here we determined a 1.92-Å resolution cryo-electron microscopic structure of a PSI-LHCI supercomplex isolated from the red alga *Cyanidium caldarium* RK-1 (NIES-2137) which is an important taxon in the Cyanidiophyceae, and subsequently investigated these correlations through structural comparisons and phylogenetic analysis. The PSI-LHCI structure shows five LHCI subunits together with a PSI-monomer core. The five LHCIs are composed of two Lhcr1s, two Lhcr2s, and one Lhcr3. Phylogenetic analysis of LHCs bound to PSI in red-lineage algae showed clear orthology of LHCs between *C. caldarium* and *Cyanidioschyzon merolae*, whereas no orthologous relationships were found between *C. caldarium* Lhcr1–3 and LHCs in other red-lineage PSI-LHCI structures. These findings provide evolutionary insights into conservation and diversity of red-lineage LHCs associated with PSI.

---

Oxygenic photosynthesis of cyanobacteria, algae, and land plants converts solar energy into chemical energy concomitant with the evolution of oxygen molecules^1^. The light-energy conversion takes place in photosystem I and photosystem II (PSI and PSII, respectively), two multi-subunit pigment-protein complexes that perform light harvesting, charge separation, and electron transfer reactions^1^. In addition to PSI and PSII, light-harvesting complexes (LHCs) participate in the acquisition of sunlight and transfer of excitation energy to PSI and PSII. PSI and PSII are thus associated with LHCI and LHCII, respectively, to form PSI-LHCI and PSII-LHCII supercomplexes^1^.

LHCs are highly diversified among photosynthetic organisms in terms of the protein sequences and pigment compositions of chlorophylls (Chls) and carotenoids (Cars) bound to the LHC proteins^1–3^. The differences of LHCs cause color variations in photosynthetic organisms, which can be classified into green and red lineages^4^. The green lineage includes green algae and land plants, whereas the red lineage includes red algae, diatoms, haptophytes, cryptophytes, and dinoflagellates^4^. The structures of LHCs and their association patterns with the photosystem cores have been revealed by structural studies, especially using cryo-electron microscopy (cryo-EM)^3,5^. In the red lineage, the number, sequences, and pigment compositions of LHCIs have been found to differ greatly among the PSI-LHCI structures of red algae^6–8^, a diatom^9,10^, and a cryptophyte^11^.

Red algae are grouped into a distinctive photosynthetic lineage including unicellular and large multicellular taxa^12^, and represent an evolutionary intermediate between cyanobacteria and red-lineage algae^4^. Two types of PSI-LHCI structures from different red algae have been reported. The red alga *Cyanidioschyzon merolae* belongs to the Cyanidiophyceae (Cyanidiophytina)^13–16^, and its PSI-LHCI structure contained three to five LHCI subunits^6,7^. In contrast, the red alga *Porphyridium purpureum* belongs to the Porphyridiophyceae^16,17^, and its PSI-LHCI structure contained seven LHCI subunits and one red lineage chlorophyll *a*/*b*-binding-like protein (RedCAP)^8^ which is included in the LHC protein superfamily^18,19^. Thus, the number and binding sites of LHCIs in PSI-LHCI differ significantly between the two types of red algae.

*Cyanidium caldarium* is a unicellular red alga belonging to the Cyanidiophyceae^13–16^ and lives in thermo-acidic environments^20^. Isolation and characterization of *C. caldarium* PSI-LHCI supercomplexes have been reported^21,22^. Gardian et al. suggested that the *C. caldarium* PSI-LHCI has 0–8 LHCIs per monomeric PSI core in response to growth-light conditions^21^. However, the overall structure of the *C. caldarium* PSI-LHCI is still unknown.

In this study, we solved a 1.92-Å resolution structure of a PSI-LHCI supercomplex purified from *C. caldarium* by single-particle cryo-EM. The structure shows a PSI-monomer core and five LHCI subunits. The arrangement of LHCIs in the *C. caldarium* PSI-LHCI structure was very similar to that of the *C. merolae* PSI-LHCI structure. Based on the structural comparisons together with phylogenetic analysis of the red-lineage LHCs bound to PSI, we discuss the molecular evolution of LHCs from red algae to diatoms and cryptophytes.

## Results and discussion

### Overall structure of the *C. caldarium* PSI-LHCI supercomplex

The PSI-LHCI supercomplex was purified from *C. caldarium* RK-1 (NIES-2137), and characterized in our previous study^22^. Cryo-EM images of the PSI-LHCI supercomplex were obtained by a JEOL CRYO ARM 300 electron microscope operated at 300 kV. The final cryo-EM map was determined with a C1 symmetry at a resolution of 1.92 Å (Supplementary Fig. 1, 2, Supplementary Table 1), based on the “gold standard” Fourier shell correlation (FSC) = 0.143 criterion (Supplementary Fig. 2a).

The atomic model of PSI-LHCI was built based on the resultant cryo-EM map (see Methods; Supplementary Fig. 2, Supplementary Table 1–3). The structure reveals a monomeric PSI core associated with five LHCI subunits (Fig. 1a, b). Three of the five LHCI subunits are located near PsaA, PsaJ, and PsaF, whereas the remaining two subunits are positioned near PsaB, PsaI, PsaL, and PsaM (Fig. 1b). The five LHCIs were named LHCI-1 to 5 (Fig. 1a). The PSI core contains 97 Chls *a*, 20 *β*-carotenes (BCRs), 2 zeaxanthins (ZXTs), 3 [4Fe-4S] clusters, 2 phylloquinones, and 4 lipid molecules, whereas the LHCI subunits contain 57 Chls *a* and 21 ZXTs (Supplementary Table 3).

**Fig. 1.**
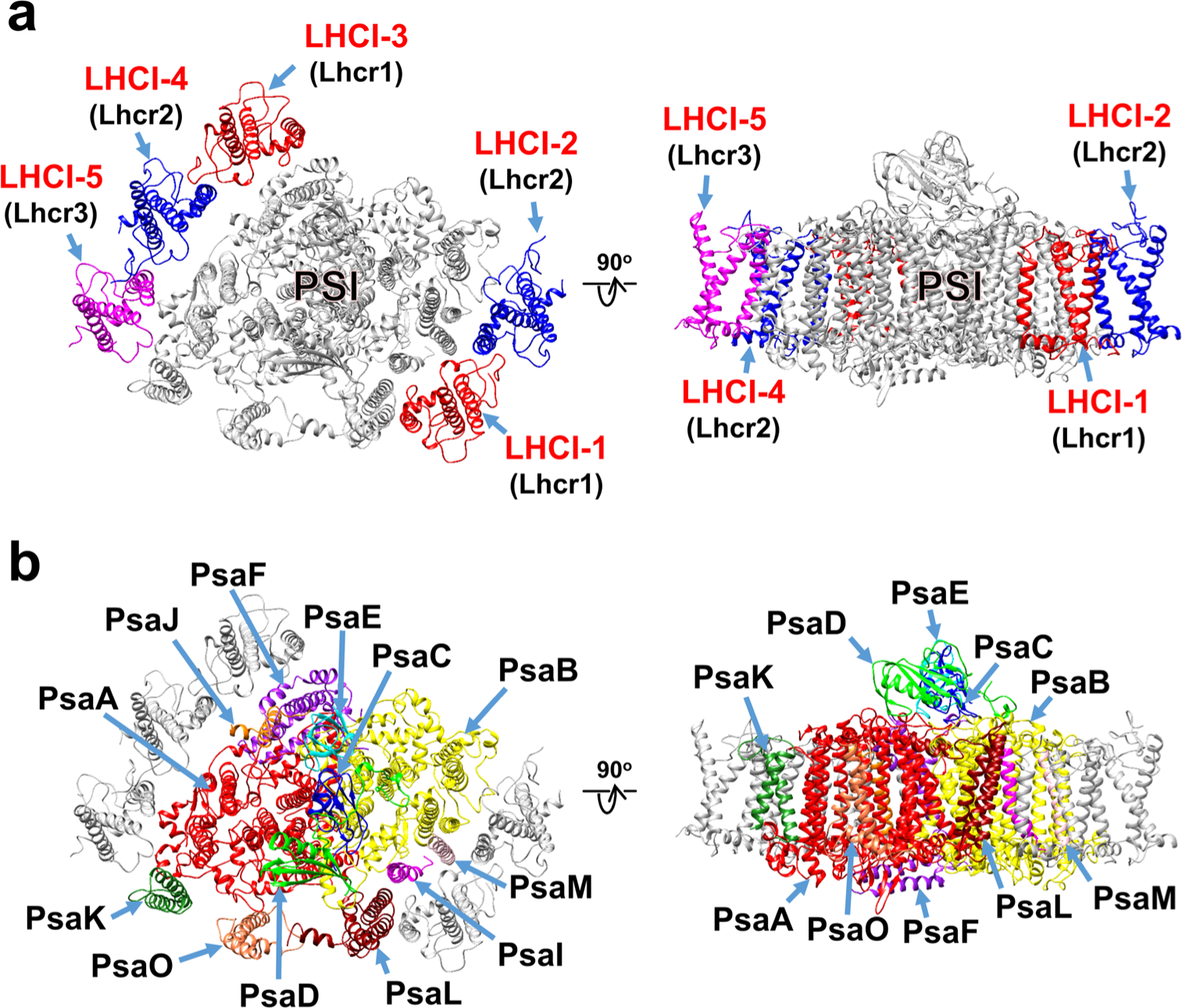
Overall structure of the PSI-LHCI supercomplex from *C. caldarium* RK-1 (NIES-2137). Structures are viewed from the stromal side (left panels) and the direction perpendicular to the membrane normal (right panels). Only protein structures are shown, and cofactors are omitted for clarity. The LHCI (**a**) and PSI-core (**b**) subunits were labeled and colored differently. **a,** The five LHCI subunits were labeled as LHCI-1 to 5 (red) with their gene products indicated in parentheses (black).

### Structure of the *C. caldarium* PSI core

The PSI-monomer core has 12 subunits, PsaA, PsaB, PsaC, PsaD, PsaE, PsaF, PsaI, PsaJ, PsaK, PsaL, PsaM, and PsaO (Fig. 1b), and their arrangements are similar to that in the *C. merolae* PSI-LHCI^6,7^. The sequences of each gene of the PSI subunits were based on the *de novo* assembly of the RNA sequencing data of *C. caldarium* RK-1 (NIES-2137). Among the PSI subunit genes, two types of PsaO, PsaO-1 and PsaO-2, were found; however, we cannot distinguish between the two PsaOs in the PSI-LHCI structure. This is because the structure of PsaO was modeled from Y64 to Y146 (Supplementary Fig. 3a), the sequences of which are the same between PsaO-1 and PsaO-2 (Supplementary Fig. 3b).

The number and arrangement of Chls in the *C. caldarium* PSI-LHCI (Supplementary Fig. 3c) are virtually identical to those in the *C. merolae* PSI-LHCI^7^. In contrast, BCR852 of PsaB in the *C. caldarium* PSI-LHCI (Supplementary Fig. 3d) was not found in the *C. merolae* PSI-LHCI^7^. ZXT304 of PsaF and ZXT105 of PsaJ were identified according to our criterion (Supplementary Fig. 3e; see Methods), and they correspond to BCR304 of PsaF and BCR105 of PsaJ, respectively, in the *C. merolae* PSI-LHCI^7^.

ZXT has a OH group in each end ring, whereas BCR does not have any OH groups in either of the rings. When we look at the surrounding structures of the end rings of PsaF-ZXT304 and PsaJ-ZXT105, no characteristic interactions are found, such as hydrogen bonds to the OH groups of ZXTs (Supplementary Fig. 3f, g). Since the protein and cofactor structures around the end rings of PsaF-ZXT304 and PsaJ-ZXT105 in the *C. caldarium* PSI-LHCI are similar to those of PsaF-BCR304 and PsaJ-BCR105 in the *C. merolae* PSI-LHCI^7^, it is suggested that the corresponding Cars are exchangeable between ZXT and BCR. However, it should be noted that the two BCRs of the *C. merolae* PSI-LHCI may be ZXTs, because the *C. merolae* PSI-LHCI structures were reported at low resolutions (3.6–4.0 Å)^6,7^, which could lead to a mis-assignment of the Cars. Further improvement of the *C. merolae* PSI-LHCI structure is required for identification of the Car molecules of PsaF and PsaJ, in order to gain structural insights into an exchangeable mechanism of Cars in photosynthetic pigment-protein complexes.

### Structure of the *C. caldarium* LHCIs

The five LHCI subunits in the PSI-LHCI structure were identified using three Lhcr genes, so that LHCI-1 and LHCI-3 were Lhcr1; LHCI-2 and LHCI-4 were Lhcr2; LHCI-5 was Lhcr3 (Fig. 1a). These arrangements correspond to those in the *C. merolae* PSI-LHCI^7^. Two types of Lhcr1, Lhcr1-1 and Lhcr1-2, were found by *de novo* assembly of the RNA sequencing data of *C. caldarium*. Similarly, Lhcr2 and Lhcr3 were also found as Lhcr2-1/Lhcr2-2 and Lhcr3-1/Lhcr3-2, respectively. The structures of LHCI-1 and LHCI-3 were modeled from Q41 to P212 (Supplementary Fig. 4a, b); in this region, the sequence of Lhcr1-1 is consistent with that of Lhcr1-2 (Supplementary Fig. 4c). The structures of LHCI-2 and LHCI-4 were modeled from Q34 to F216 (Supplementary Fig. 4d, e). The 40th amino acid of Lhcr2-1 is alanine, whereas that of Lhcr2-2 is threonine (Supplementary Fig. 4f), and the 40th amino acid of LHCI-2 and LHCI-4 was provisionally modeled as alanine. The structure of LHCI-5 was modeled from A63 to V213 (Supplementary Fig. 4g); in this region, the sequence of Lhcr3-1 is consistent with that of Lhcr3-2 (Supplementary Fig. 4h).

For the assignments of each LHCI subunit, we focused on characteristic amino-acid residues among the three types of Lhcrs. As for LHCI-1 and LHCI-3, Lhcr1 was identified because its amino-acid residues Y61 and R62 are different from the corresponding residues in Lhcr2 and Lhcr3 (Supplementary Fig. 5a, b, d). As for LHCI-2 and LHCI-4, Lhcr2 was identified because its amino-acid residues G68 and F69 are different from the corresponding residues in Lhcr1 and Lhcr3 (Supplementary Fig. 5a, c, e). As for LHCI-5, Lhcr3 was identified because its amino-acid residues Y73 and L74 are different from the corresponding residues in Lhcr1 and Lhcr2 (Supplementary Fig. 5a, f).

LHCI-1 binds 9 Chls *a* and 4 ZXTs (Fig. 2a). The axial ligands of the central Mg atoms of Chls within LHCI-1 are mainly provided by main and side chains of amino acid residues (Supplementary Table 4). In contrast, LHCI-3 contains 11 Chls *a* and 5 ZXTs (Fig. 2c). The axial ligands of the LHCI-3 Chls are summarized in Supplementary Table 4. The number of Chls and ZXTs in LHCI-1 is less than those in LHCI-3, albeit with the same gene product between the two subunits. This may be due to weaker densities for Chls and ZXTs in LHCI-1 than those in LHCI-3, leading to the inability of pigment assignment according to our criterion (see Methods). Alternatively, it may be possible that some Chls may originally be absent in LHCI-1. Because LHCI-1 and LHCI-3 are located near PsaB and PsaA, respectively (Fig. 1), the different pigment content between LHCI-1 and LHCI-3 may result in variations in excitation-energy transfer from LHCIs to PSI and/or among LHCIs.

**Fig. 2.**
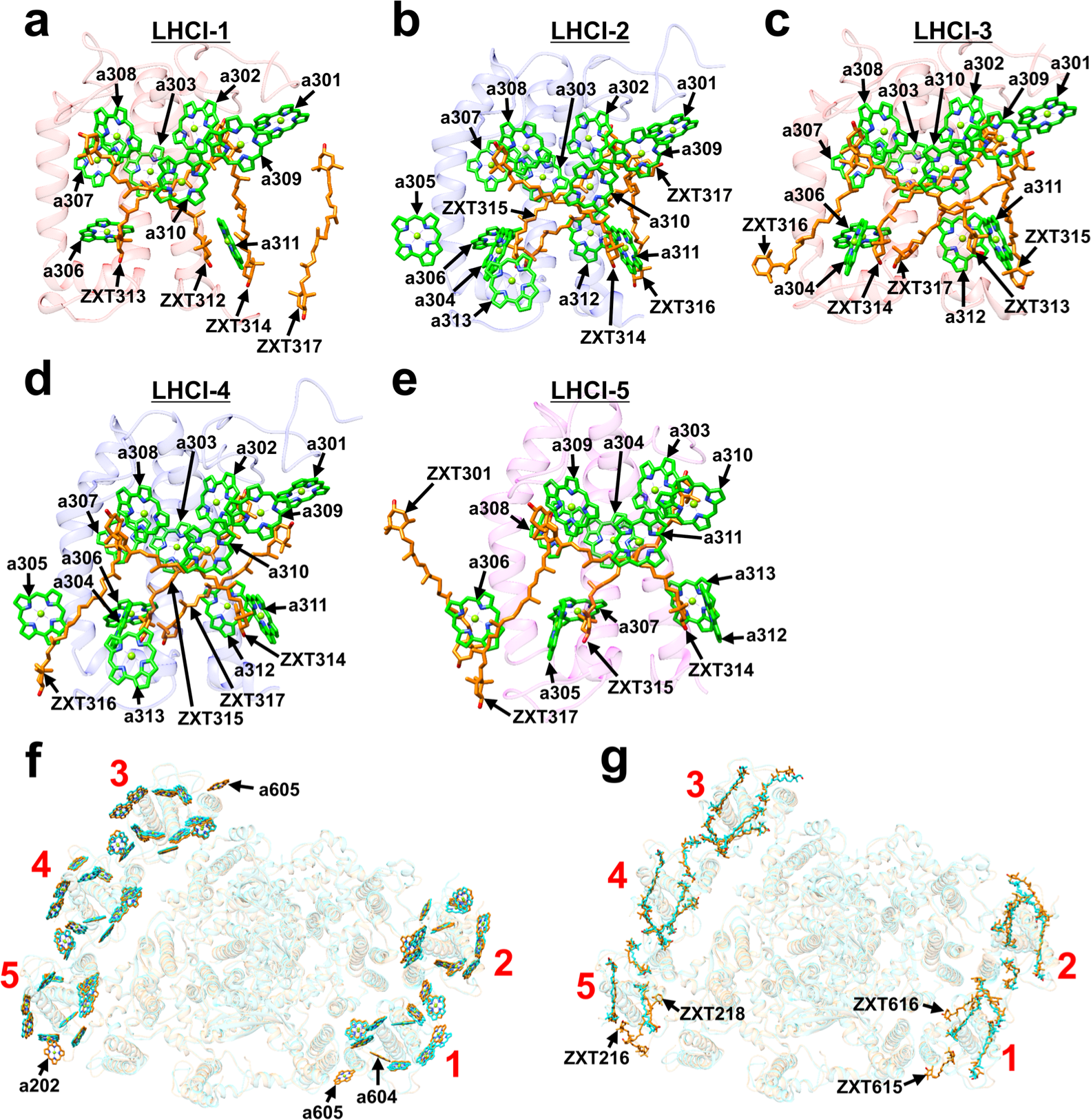
Structure of LHCIs. **a–e,** Structures of LHCI-1 to LHCI-5 depicted as transparent cartoon models and arrangements of Chl and ZXT. Chls and ZXTs are colored green and orange, respectively. Only rings of the Chl molecules are depicted. **f, g,** Comparison of Chls (**f**) and Cars (**g**) of PSI-LHCI between *C. caldarium* (cyan) and *C. merolae* (orange). The *C. caldarium* PSI-LHCI structure was superimposed with the *C. merolae* PSI-LHCI structure (PDB: 5ZGB). The structures are viewed from the stromal side. Chls and Cars are shown as sticks. Only rings of the Chl molecules are depicted. Numbers 1 to 5 (red) indicate the sites of LHCI-1 to 5, respectively, in the *C. caldarium* PSI-LHCI structure.

LHCI-2 and LHCI-4 each bind 13 Chls *a* and 4 ZXTs (Fig. 2b, d), whereas LHCI-5 binds 11 Chls *a* and 4 ZXTs (Fig. 2e). The Chl ligands of LHCI-2, LHCI-4, and LHCI-5 are mainly amino acid residues (Supplementary Table 4). The root mean square deviations (RMSDs) of the structures between LHCI-3 and the other four LHCIs range from 0.74 to 1.55 Å for 302 Cα atoms (Supplementary Table 5).

The arrangement of Chls and Cars in the LHCI subunits of *C. caldarium* PSI-LHCI is virtually identical to those in the *C. merolae* PSI-LHCI^7^ (Fig. 2f, g); however, their numbers are slightly fewer in the *C. caldarium* PSI-LHCI than those in the *C. merolae* PSI-LHCI^7^ (Fig. 2f, g). The pigment molecules of a604/a605/ZXT615/ZXT616 at the LHCI-1 site, a605 at the LHCI-3 site, and a202/ZXT216/ZXT218 at the LHCI-5 site were observed in the *C. merolae* PSI-LHCI structure but not in the *C. caldarium* PSI-LHCI structure (Fig. 2f, g). These results imply different excitation-energy-transfer mechanisms of LHCIs between *C. caldarium* and *C. merolae*.

### Structural comparisons of the *C. caldarium* LHCIs with those of other red algae

We compared binding sites of LHCIs in the PSI-LHCI structures between *C. caldarium* and *C. merolae* (Fig. 3a). The five LHCI subunits of LHCI-1 to LHCI-5 in the *C. caldarium* PSI-LHCI structure (red) are located at the same positions of Lhcr1*, Lhcr2*, Lhcr1, Lhcr2, and Lhcr3, respectively, in the *C. merolae* PSI-LHCI structure (cyan)^7^ (Fig. 3a). The gene products of each LHC subunit are shown in Fig. 3b, and multiple sequence alignments of the LHC proteins located at the same positions in the PSI-LHCI structures of *C. caldarium* and *C. merolae* were shown in Supplementary Fig. 6a–c. The *C. caldarium* Lhcr1–3 show high sequence similarities (89–91%) to the *C. merolae* Lhcr1– 3, respectively. These results provide evidence for strong conservation of the binding sites of individual LHCIs to PSI and their genes between *C. caldarium* and *C. merolae*. It should be pointed out that in *C. merolae*, two types of PSI-LHCI have been reported, namely, one has three LHCI subunits associated at one side of the PSI core whereas the other one has five LHCI subunits^7^. In *C. caldarium*, we only observed the five LHCIs-type of PSI-LHCI but the three LHCIs-type particles were not observed. This may be due to a low light intensity for the growth of the *C. caldarium* cells, or alternatively, PSI-LHCI exists as the five LHCIs-type only in *C. caldarium*.

**Fig. 3.**
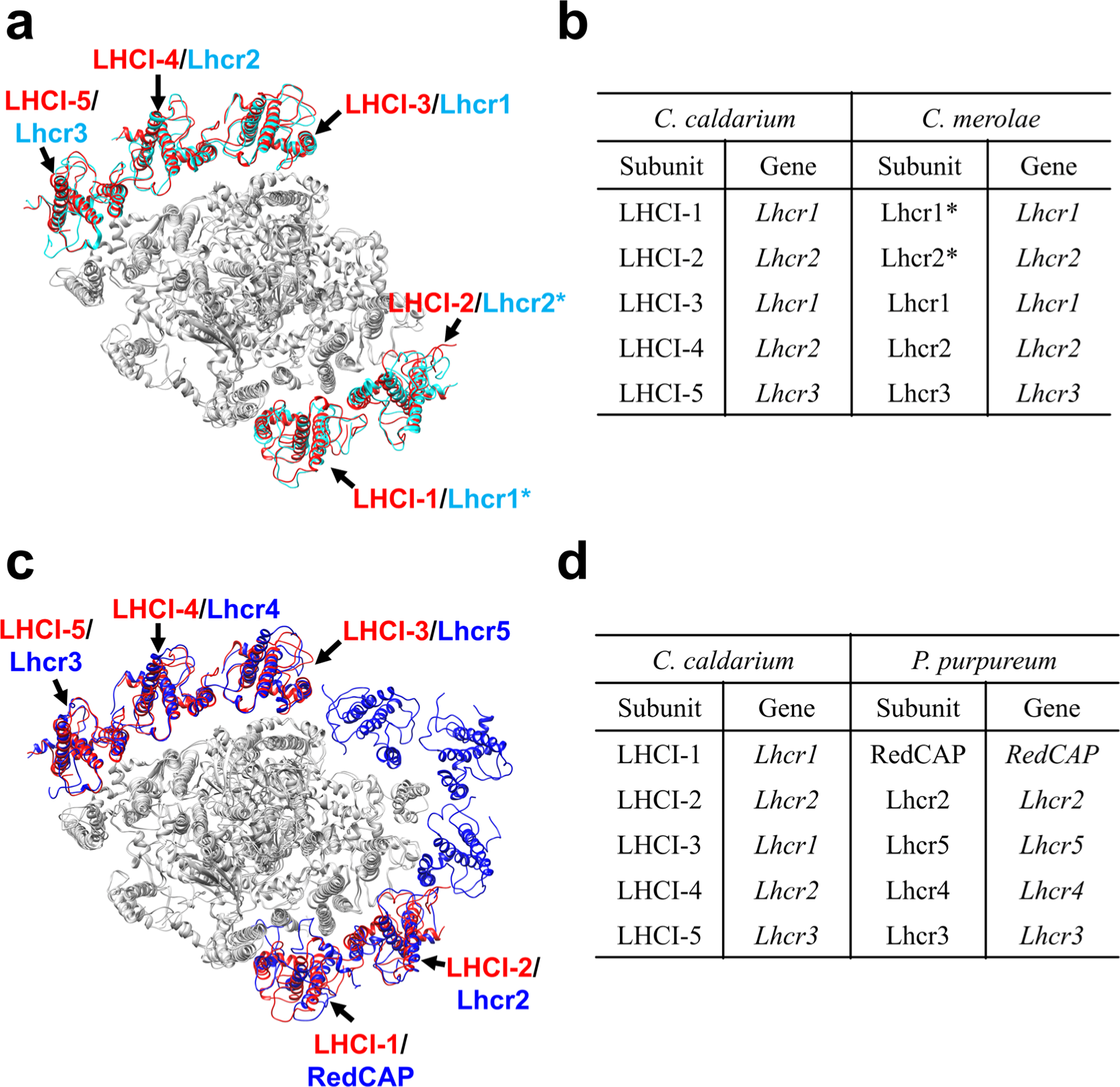
Structural comparisons of the *C. caldarium* LHCIs with those of other red algae. **a,** Superposition of the *C. caldarium* PSI-LHCI structure with the *C. merolae* PSI-LHCI structure (PDB: 5ZGB). The LHCI subunits of *C. caldarium* and *C. merolae* were colored red and cyan, respectively. The structures are viewed from the stromal side. **b,** Correlation of the names of LHCs in the PSI-LHCI structures with their genes between *C. caldarium* and *C. merolae*. The names of LHCs and their genes are derived from Pi et al. (2018) for *C. merolae*^7^. **c,** Superposition of the *C. caldarium* PSI-LHCI structure with the *P. purpureum* PSI-LHCI structure (PDB: 7Y5E). The LHCI subunits of *C. caldarium* and *P. purpureum* were colored red and blue, respectively. The structures are viewed from the stromal side. **d,** Correlation of the names of LHCs in the structures with their genes between *C. caldarium* and *P. purpureum*. The names of LHCs and their genes are derived from You et al. (2023) for *P. purpureum*^8^.

The *C. caldarium* LHCIs of LHCI-1 to LHCI-5 (red) exist at the same positions of RedCAP, Lhcr2, Lhcr5, Lhcr4, and Lhcr3, respectively, in the *P. purpureum* PSI-LHCI structure (blue)^8^ (Fig. 3c). The gene products of each LHC subunit are shown in Fig. 3d, and multiple sequence alignments of the LHC proteins located at the same positions in the PSI-LHCI structures of *C. caldarium* and *P. purpureum* were shown in Supplementary Fig. 7a–e. The amino acid sequence of *C. caldarium* Lhcr1 has a low similarity of 31% with that of *P. purpureum* RedCAP. Since RedCAPs are one of the LHC protein superfamily but are different from the LHC protein family^18,19^, the binding site at LHCI-1 in the *C. caldarium* PSI-LHCI structure may be diversified between the Cyanidiophyceae and Porphyridiophyceae. In contrast, sequence alignments between *C. caldarium* Lhcr2 and *P. purpureum* Lhcr2, between *C. caldarium* Lhcr1 and *P. purpureum* Lhcr5, between *C. caldarium* Lhcr2 and *P. purpureum* Lhcr4, and between *C. caldarium* Lhcr3 and *P. purpureum* Lhcr3 show similarities of 52–64%, which supports their occupations of the same sites in the different PSI-LHCIs.

### Structural comparison of the *C. caldarium* LHCIs with the diatom *Chaetoceros gracilis* FCPIs

Diatoms have unique LHCs, fucoxanthin Chl *a*/*c*-binding proteins (FCPs), whose bindings to PSI have been shown by cryo-EM single-particle analysis^9,10^. Here, we compared binding sites of the *C. caldarium* LHCIs with the *Chaetoceros gracilis* FCPIs^9^ (Fig. 4a). The four LHCI subunits, LHCI-2 to LHCI-5, in the *C. caldarium* PSI-LHCI structure (red) are located at the same positions as Fcpa1, Fcpa5, Fcpa6, and Fcpa7, respectively, in the *C. gracilis* PSI-FCPI structure (orange)^9^, whereas the *C. caldarium* LHCI-1 site is empty in the *C. gracilis* PSI-FCPI structure (Fig. 4a). The gene products of each LHC subunit are shown in Fig. 4b, and multiple sequence alignments were generated for LHC proteins located at the same positions in the *C. caldarium* PSI-LHCI and *C. gracilis* PSI-FCPI structures (Supplementary Fig. 8a–d). The results of sequence alignments between *C. caldarium* Lhcr2 and *C. gracilis* Lhcr1, between *C. caldarium* Lhcr1 and *C. gracilis* Lhcr5, between *C. caldarium* Lhcr2 and *C. gracilis* Lhcr6, and between *C. caldarium* Lhcr3 and *C. gracilis* Lhcr7 show similarities in the range of 45– 50%.

**Fig. 4.**
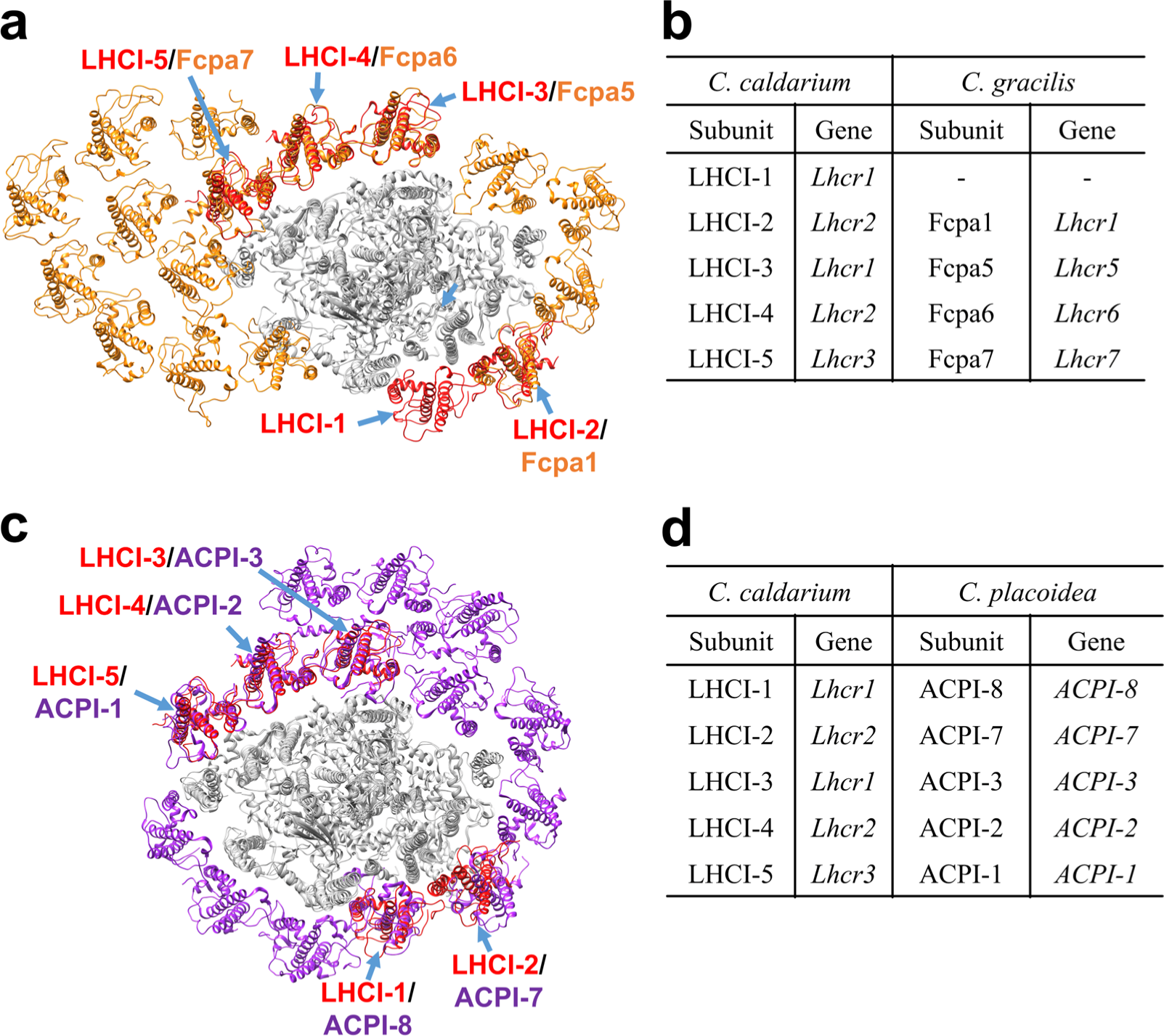
Structural comparisons of the *C. caldarium* LHCIs with the diatom *C. gracilis* FCPIs and the cryptophyte *C. placoidea* ACPIs. **a,** Superposition of the *C. caldarium* PSI-LHCI structure with the *C. gracilis* PSI-FCPI structure (PDB: 6L4U). The *C. caldarium* LHCI and *C. gracilis* FCPI subunits were colored red and orange, respectively. The structures are viewed from the stromal side. **b,** Correlation of the names of LHCs in the structures with their genes between *C. caldarium* and *C. gracilis*. The names of LHCs and their genes are derived from Nagao et al. (2020) for *C. gracilis*^9^. **c,** Superposition of the *C. caldarium* PSI-LHCI structure with the *C. placoidea* PSI-ACPI structure (PDB: 7Y7B). The *C. caldarium* LHCI and *C. placoidea* ACPI subunits were colored red and purple, respectively. The structures are viewed from the stromal side. **d,** Correlation of the names of LHCs in the structures with their genes between *C. caldarium* and *C. placoidea*. The names of LHCs and their genes are derived from Zhao et al. (2023) for *C. placoidea*^11^.

It is known that antenna sizes of FCPIs in the *C. gracilis* PSI-FCPI supercomplexes are altered in response to growth conditions, especially CO_2_ concentrations and temperatures^23^. This has been shown by the presence of two different PSI-FCPI structures, one with 16 FCPI subunits^9^ and the other with 24 FCPI subunits^10^ (Supplementary Fig. 9a). Different from our PSI-FCPI structure with 16 FCPIs^9^, each FCP subunit in the PSI-FCPI structures with 24 FCPI subunits^10^ was identified using transcriptome data of *C. gracilis* as well as FCP sequences from other diatom species. The amino acid sequences of 16 FCPIs observed in our PSI-FCPI structure^9^ were different from those in the PSI-FCPI structures with 24 FCPI subunits^10^. Moreover, the gene names of the 16 FCPIs differ between our structure^9^ and the other^10^, which were discussed previously^10,24^. As our PSI-FCPI structure^9^ was built according to the genome data of *C. gracilis*^24^, we performed further data analysis and discussion of the *C. caldarium* LHCI-2 to LHCI-5 using the structural data of our PSI-FCPI supercomplex^9^ (Fig. 4a,b) in the present study.

It is particularly noteworthy that the PSI-FCPI structure with 24 FCPIs showed a FCPI subunit located at the *C. caldarium* LHCI-1 site (Supplementary Fig. 9a). The FCPI subunit was built using an FCP sequence of a diatom *Fragilariopsis cylindrus*, which was *fc13194* (Supplementary Fig. 9b). This gene is a RedCAP as discussed in Kumazawa et al. 2022^24^. Here we searched this gene in the *C. gracilis* database, and obtained a gene like RedCAP in *C. gracilis*, which was termed as *CgRedCAP*. A sequence alignment between fc13194 and CgRedCAP (Supplementary Fig. 9c) shows a similarity of 50%. Thus, the FCPI subunit located at the same position of the *C. caldarium* LHCI-1 may be a RedCAP (Gene ID: g6493.t1) in *C. gracilis*.

### Structural comparison of the *C. caldarium* LHCIs with the cryptophyte *Chroomonas placoidea* ACPIs

Cryptophytes have unique LHCs, alloxanthin Chl *a*/*c*-binding proteins (ACPs), whose bindings to PSI have been shown as a PSI-ACPI supercomplex by cryo-EM single-particle analysis^11^. We compared binding sites of the *C. caldarium* LHCIs with the *Chroomonas placoidea* ACPIs (Fig. 4c). The *C. placoidea* PSI-ACPI structure showed 14 ACPI subunits in total, which is much more than LHCIs found in the red algae. The five LHCI subunits of LHCI-1 to LHCI-5 in the *C. caldarium* PSI-LHCI structure (red) are located at the same positions of ACPI-8, ACPI-7, ACPI-3, ACPI-2, and ACPI-1, respectively, in the *C. placoidea* PSI-ACPI structure (purple)^11^ (Fig. 4c). The gene products of each LHC subunit are shown in Fig. 4d, and multiple sequence alignments were generated for the LHC proteins located at the same positions in the *C. caldarium* PSI-LHCI and *C. placoidea* PSI-ACPI structures (Supplementary Fig. 10a–e). The results of sequence alignments between *C. caldarium* Lhcr2 and *C. placoidea* ACPI-7, between *C. caldarium* Lhcr1 and *C. placoidea* ACPI-3, between *C. caldarium* Lhcr2 and *C. placoidea* ACPI-2, and between *C. caldarium* Lhcr3 and *C. placoidea* ACPI-1 show similarities of 49–59%. As for ACPI-8, sequence alignment between *C. caldarium* Lhcr1 and *C. placoidea* ACPI-8 shows a low similarity of 37%. Compared with RedCAPs of *P. purpureum* and *C. gracilis*, ACPI-8 has sequence similarities of 42 and 56%, respectively (Supplementary Fig. 11), indicating that ACPI-8 is a RedCAP.

### Phylogenetic relationships among the structurally known red-lineage LHCs

We further examined evolutionary relationships of the *C. caldarium* LHCI-1 to LHCI-5 with other red-lineage LHCs in the PSI-LHCI structures including PSI-FCPI and PSI-ACPI by phylogenetic analysis (Fig. 5a). At the site of *C. caldarium* LHCI-1, the *C. caldarium* Lhcr1 (CcLhcr1) and *C. merolae* Lhcr1 (CmLhcr1) belonged to the same clade in the phylogenetic tree (Fig. 5a), whereas *P. purpureum*, *C. gracilis*, and *C. placoidea* have no Lhcr but RedCAP (Fig. 5b). Since RedCAPs were phylogenetically distinct from the LHC protein family but assumed to serve as a light harvesting antenna and belonged to the LHC protein superfamily^18,19^, the LHC subunits at the *C. caldarium* LHCI-1 site can be divided into two groups: *C. caldarium*/*C. merolae* Lhcr1s and *P. purpureum*/*C. gracilis*/*C. placoidea* RedCAPs.

**Fig. 5.**
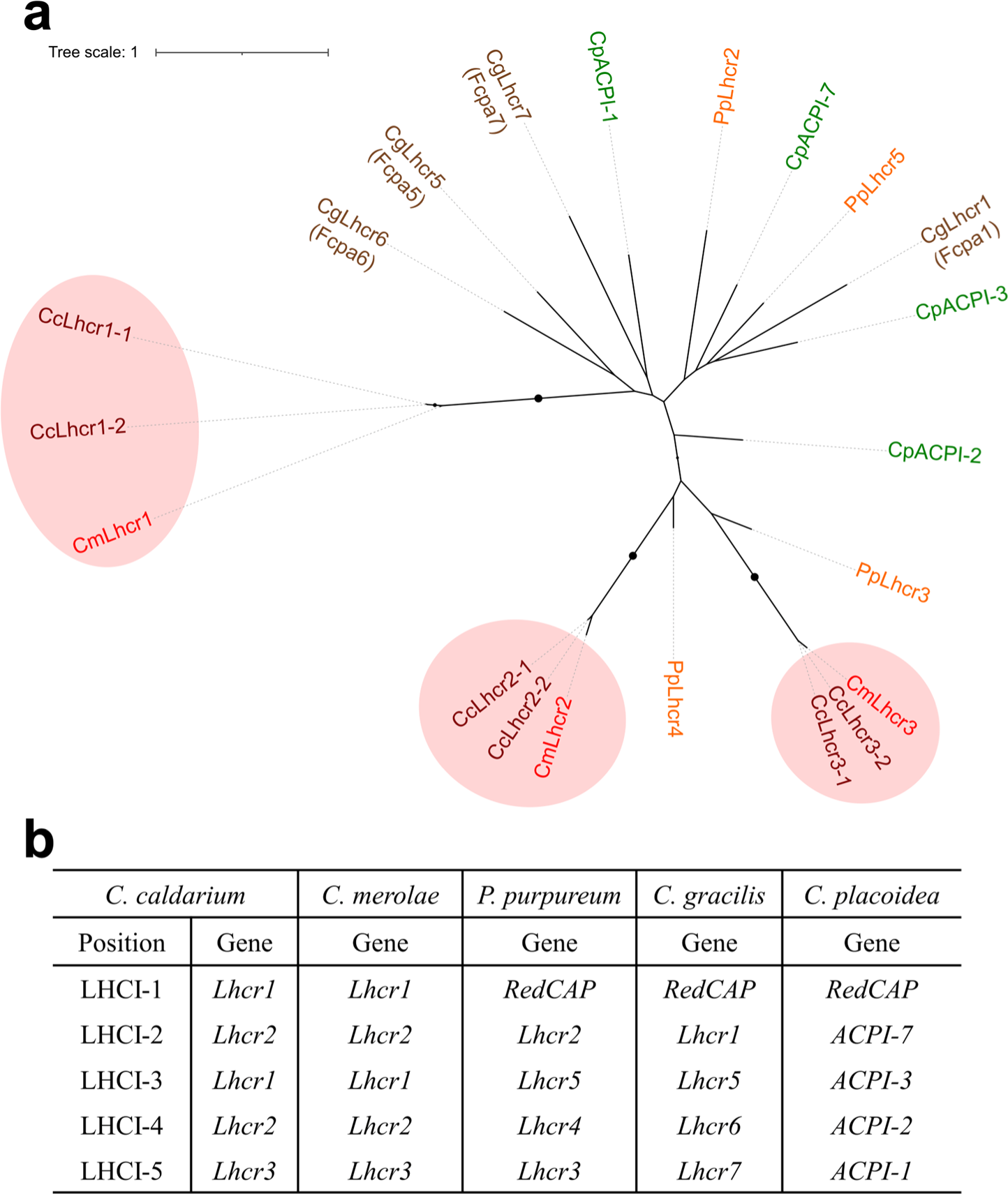
Phylogenetic analysis and correlation of LHCs among red-lineage algae. **a,** Unrooted maximum-likelihood tree of the LHC proteins whose binding sites to PSI are conserved between *C. caldarium* and other red-lineage algae, except for RedCAPs. The phylogenetic tree was inferred by IQ-TREE 2 using the LG+I+G4 model and the trimmed alignment of 21 sequences with 268 amino-acid residues. Black circles at branches indicate ≥95% ultrafast bootstrap support (1000 replicates). The names of red-lineage algae were written in front of each gene name; for example, CcLhcr1-2 means Lhcr1-2 of *C. caldarium*. Cm, *C. merolae*; Pp, *P. purpureum*; Cg, *C. gracilis*; Cp, *C. placoidea*. **b**, Correlation of the genes of LHCs in the structures with their binding positions based on the *C. caldarium* LHCI-1 to 5. Each gene was explained in Fig. 3, 4, and Supplementary Fig. 9; however, RedCAPs of *C. gracilis* and *C. placoidea* were interpreted in the present study.

At the site of *C. caldarium* LHCI-2, CcLhcr2 and CmLhcr2 belonged to the same clade in the phylogenetic tree, but Lhcr2 of *P. purpureum* (PpLhcr2), Lhcr1 of *C. gracilis* (CgLhcr1), and ACPI-7 of *C. placoidea* (CpACPI-7) did not exist in the same clade of CcLhcr2 (Fig. 5a, b). In addition, PpLhcr5, CgLhcr5, and CpACPI-3 were not included in the same clade of CcLhcr1 at the site of *C. caldarium* LHCI-3, whereas PpLhcr4, CgLhcr6, and CpACPI-2 were not grouped into the same clade of CcLhcr2 at the site of *C. caldarium* LHCI-4 (Fig. 5a, b). At the site of *C. caldarium* LHCI-5, CcLhcr3 and CmLhcr3 belonged to the same clade in the phylogenetic tree, but PpLhcr3, CgLhcr7, and CpACPI-1 did not exist in the same clade of CcLhcr3 (Fig. 5a, b). Thus, our phylogenetic tree strongly indicates that the *C. caldarium* Lhcr1, Lhcr2, and Lhcr3 are orthologous to the *C. merolae* Lhcr1, Lhcr2, and Lhcr3, respectively, at each site of LHCI-1 to 5 in the *C. caldarium* PSI-LHCI structure. However, the orthologous relationships of CcLhcr1–3 with LHCs in the *P. purpureum* PSI-LHCI, *C. gracilis* PSI-FCPI, and *C. placoidea* PSI-ACPI structures could not be supported at any sites of the *C. caldarium* LHCI-1 to 5 in our phylogenetic tree.

### Evolution of the red-lineage LHCs bound to PSI

Structural comparisons of LHCs in the *C. caldarium* PSI-LHCI structure with the structures of other red-lineage algae reveal conservation of the binding positions of *C. caldarium* LHCI-1 to LHCI-5 (Fig. 3, 4, Supplementary Fig. 9). In contrast, our phylogenetic analysis showed conservation and diversity of the structurally known LHCs among the red-lineage algae (Fig. 5), even though both diatoms and cryptophytes are thought to evolve from red algae^4^. It is known that RedCAPs are grouped into one of the ancestral LHC protein family in the red lineage^18,19^ and they appear to be already provided in an ancestral red alga prior to the divergence between Rhodophytina and the monophyletic Cyanidiophyceae including the order Cyanidiales, Cyanidioschyzonales, and Galdieriales^16,18,19,25^. *Galdieria sulphuraria* is classified into the order Galdieriales and retains RedCAP in its genome, whereas no RedCAPs have been found at least in the genome of *C. merolae*^26^, which belongs to the order Cyanidioschyzonales^16^.

Based on the structural and phylogenetic findings, we propose a schematic model for evolution of red-lineage LHCs that bind to PSI (Fig. 6). An ancestral red alga appears to have RedCAP^18,19^, which has been found in the PSI-LHCI structures including PSI-FCPI and PSI-ACPI from *P. purpureum*, *C. gracilis*, and *C. placoidea*. This suggests that the binding property of RedCAP in the PSI-LHCI structures including PSI-FCPI and PSI-ASPI are conserved among the Porphyridiophyceae, diatoms, and cryptophytes, as well as an ancestral red alga. In contrast, the PSI-LHCI structures of *C. caldarium* and *C. merolae* have Lhcr1 instead of RedCAP at the LHCI-1 site (Red subunits in Fig. 6), suggesting that the two types of red algae choose Lhcr1 to associate with PSI instead of RedCAP after the divergence from the order Galdieriales. Since the classification of RedCAPs is different from that of the LHC protein family including CcLhcr1 and CmLhcr1 in the LHC protein superfamily^18,19^, it is highly possible that the *C. caldarium* and *C. merolae* PSI-LHCIs possess LHCs with completely different properties at the LHCI-1 site in the process of evolution from an ancestral red alga.

**Fig. 6.**
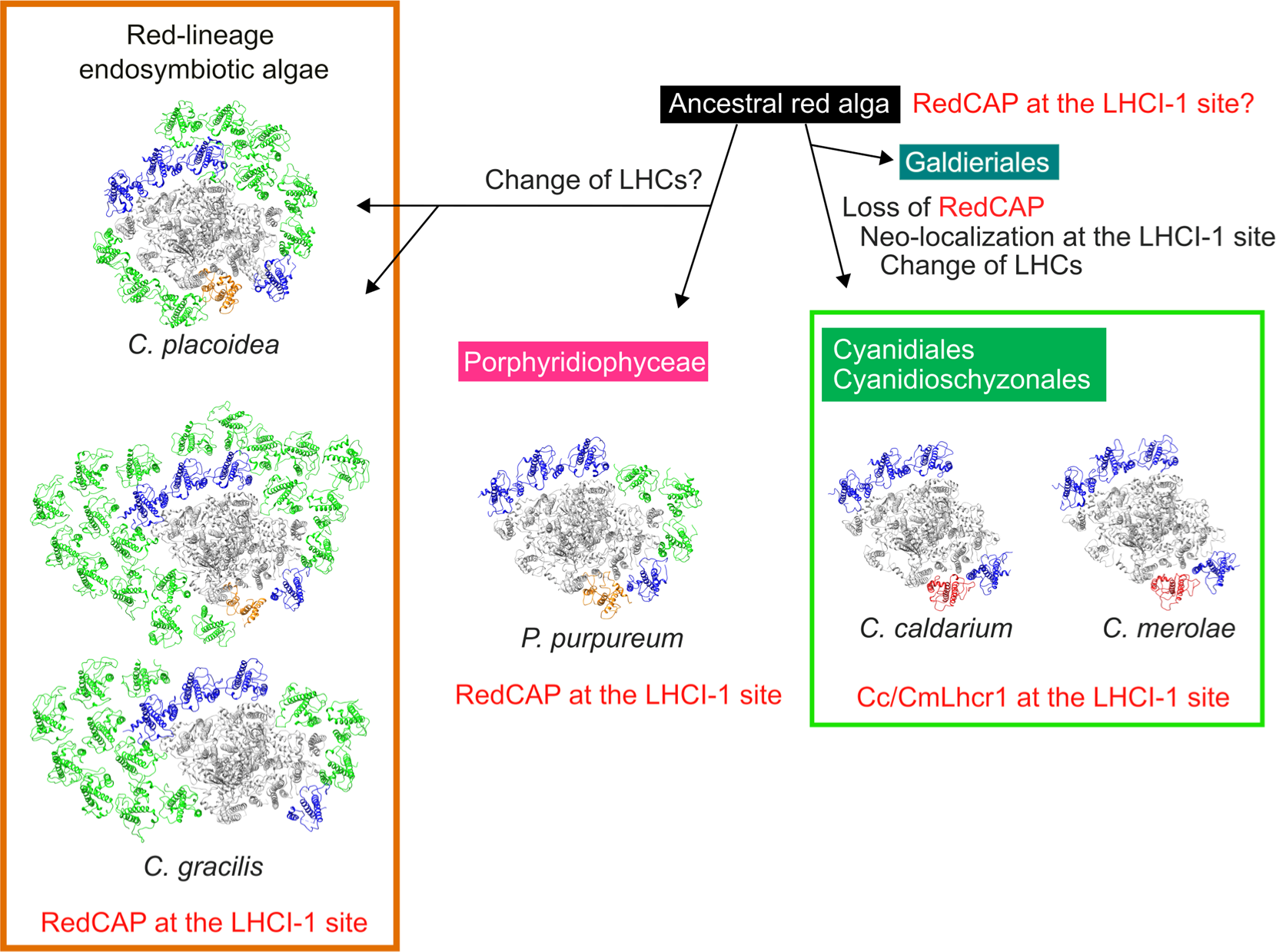
Evolutionary scheme of red-lineage LHCs in the PSI-LHCI structures. Structures are viewed from the stromal side. Grey, PSI; red and orange, the LHCI-1 site; blue, the LHCI-2 to 5 sites; green, other LHCs. The red and orange subunits indicate Lhcr1s and RedCAPs, respectively. A detailed explanation of this scheme is described in the main text.

Yoon and co-workers have shown that both the order Galdierales and Cyanidiales/Cyanidioschyzonales lose massive genes during evolution of the Cyanidiophyceae^27^. Such evolutionary adaptation may result in the loss of RedCAP and the reduced number of LHC genes after the divergence of Galdierales from other Cyanidiophyceae. At the LHCI-1 site, the space generated by the loss of RedCAP is filled by CcLhcr1 and CmLhcr1 in the *C. caldarium* and *C. merolae* PSI-LHCIs. Here we define this evolutionary event as neo-localization: a phenomenon in which a structural defect caused by a gene loss is complemented or modified by the product of another existing gene. This may have taken place at the LHCI-1 site during evolution from an ancestral red alga to the order Cyanidiales and Cyanidioschyzonales.

At the other sites of LHCI-2 to 5 (blue subunits in Fig. 6), prominent orthologous relationships were found only for CcLhcr1–3 and CmLhcr1–3 both structurally and phylogenetically. The unique LHC features of (i) clear orthology between the order Cyanidiales and Cyanidioschyzonales, (ii) neo-localization at the LHCI-1 site, and (iii) diversity of RedCAP and LHC protein families in the red lineages, provide a new evolutionary model for red-lineage LHCs. Thus, our structural and phylogenetic findings together with the evolutionary scheme of LHCs will open a new avenue for conservation and diversity of LHCs associated with PSI among red-linage algae.

## Methods

### Cryo-EM data collection

Isolation and characterization of the PSI-LHCI supercomplexes from the red alga *C. caldarium* RK-1 (NIES-2137) have been published previously^22^. For cryo-EM experiments, 3-μL aliquots of the *C. caldarium* PSI-LHCI (2.83 mg of Chl mL^−1^) in a 20 mM MES-NaOH (pH 6.5) buffer containing 5 mM CaCl_2_, 10 mM MgCl_2_, and 0.03% *β*-DDM were applied to gold pre-sputtered Quantifoil R0.6/1 Cu 200 mesh grids in the chamber of FEI Vitrobot Mark IV (Thermo Fisher Scientific). Then, the grids were blotted with a filter paper for 6 sec at 4°C under 100% humidity. The grids were plunged into liquid ethane cooled by liquid nitrogen and then transferred into a CRYO ARM 300 electron microscope (JEOL) equipped with a cold-field emission gun operated at 300 kV. AI-assisted hole detection and data collection were carried out using a combination of yoneoLocr^28^, SerialEM^29^, and JAFIS Tool version 1 (JEOL). All image stacks were collected from 5 × 5 holes per stage adjustment to the central hole and image shifts were applied to the surrounding holes while maintaining an axial coma-free condition. The images were recorded with an in-column energy filter with a slit width of 20 eV and at a nominal magnification of × 100,000 on a direct electron detector (Gatan K3, AMETEK). The nominal defocus range was −1.8 to −0.8 μm. Physical pixel size corresponded to 0.495 Å. Each image stack was exposed at a dose rate of 16.18 e^−^Å^−2^sec^−1^ for 2.46 sec in the super resolution and CDS mode with dose-fractionated 50 movie frames. In total 17,650 image stacks were collected.

### Amino acid sequences of PSI and LHCI subunits

Genomic DNA of *C. caldarium* was extracted, and sequencing libraries were prepared with a KAPA hyper prep kit (PCR free). The paired-end (800-bp) library was sequenced by MiSeq (Illumina, Inc.) with the MiSeq reagent kit version 3 (600 cycles; Illumina). The reads were cleaned up using the cutadapt ver. 3.1^30^ by trimming low-quality ends (<QV30) and adapter sequences and by removing reads shorter than 50 bp. For *de novo* assembly of whole-genome, the trimmed reads were assembled using SPAdes ver. 3.15.0^31^. A BLASTX search was performed on the draft genome data of *C. caldarium* using amino acid sequences of PSI and LHCI subunits of *C. merolae* as queries. Each nucleic acid sequences were translated to amino acid sequences by the ExPASy Translate tool.

### Cryo-EM image processing

The resultant movie frames were aligned and summed using MotionCor2^32^ to yield dose-weighted images. Estimation of the contrast transfer function (CTF) was performed using CTFFIND4^33^. All of the following processes were performed using RELION3.1^34^. In total 3,063,750 particles were automatically picked up and used for reference-free 2D classification. Then, 1,461,422 particles were selected from good 2D classes and subsequently subjected to 3D classification without any symmetry. An initial model for the first 3D classification was generated *de novo* from 2D classification. As shown in Supplementary Fig. 1c, the final PSI-LHCI structure was reconstructed from 228,449 particles. The overall resolution of the cryo-EM map was estimated to be 1.92 Å by the gold-standard FSC curve with a cut-off value of 0.143 (Supplementary Fig. 2a)^35^. Local resolutions were calculated using RELION (Supplementary Fig. 2c).

### Model building and refinement

Two types of the cryo-EM maps were used for the model building of the PSI-LHCI supercomplex: one was a postprocessed map, and the other was a denoised map using Topaz version 0.2.4^36^. The postprocessed map was denoised using the trained model in 100 epochs with two half-maps. Each subunit of the homology models constructed using the Phyre2 server^37^ was first manually fitted into the two maps using UCSF Chimera^38^, and then their structures were inspected and manually adjusted against the maps with Coot^39^. Each model was built based on interpretable features from the density maps with the contour levels of 2.5 and 2.0 σ in the denoised and postprocessed maps, respectively. Although the *C. caldarium* LHCIs were enriched in ZXTs but with minor components of BCR and *β*-cryptoxanthin^22^, the present structure of LHCI could not distinguish among Cars. Therefore, all Car molecules of LHCIs in the PSI-LHCI structure were modeled as ZXT with the above thresholds. The PSI-LHCI structure was refined with phenix.real_space_refine^40^ and Servalcat^41^ with geometric restraints for the protein-cofactor coordination. The final model was validated with MolProbity^42^, EMRinger^43^, and *Q*-score^44^. The statistics for all data collection and structure refinement are summarized in Supplementary Table 1, 2. All structural figures were made by PyMOL^45^, UCSF Chimera, and UCSF ChimeraX^46^.

Since the numbering of Chls and Cars in this paper were different from those of the PDB data, we listed the relationship of the pigment numbering in this paper with those in the PDB data in Supplementary Table 6, 7.

### Phylogenetic analysis

Amino-acid sequences of LHCs were aligned using mafft-linsi v7.490^47^. The alignment was trimmed using ClipKit v1.4.1 with smart-gap mode. The phylogenetic tree was inferred using IQ-TREE 2^48^ with the model selected by ModelFinder^49^. The tree was visualized by iTOL v6^50^. Ultrafast bootstrap approximation was performed with 1000 replicates^51^.

## Data availability

Atomic coordinate and cryo-EM maps for the reported structure have been deposited in the Protein Data Bank under an accession code 8WEY [https://www.rcsb.org/structure/8WEY] and in the Electron Microscopy Data Bank under an accession code EMD-37480 [https://www.ebi.ac.uk/emdb/EMD-37480]. Source data are provided with this paper.

## Acknowledgements

We thank Kumiyo Kato and Satoko Kakiuchi for their assistance in this study. This work was supported by JSPS KAKENHI grant Nos. JP22KJ2017 (M.K.), JP23H02347 (K.I.), JP23K14211 (Y.N.), JP22H04916 (J.-R.S.), and JP23H02423 (R.N.), JST-Mirai Program Grant Number JPMJMI20G5 (K.Y.), Sumitomo Foundation (R.N.), Takeda Science Foundation (R.N., Koji.K.), and Research Support Project for Life Science and Drug Discovery (Basis for Supporting Innovative Drug Discovery and Life Science Research (BINDS)) from AMED under Grant Number JP23ama121006 (T.H., Keisuke.K., K.Y.).

## Author Contributions

R.N. conceived the project; R.N. prepared the PSI-LHCI supercomplex; S.H., Y.H., and S.-y.M. provided the amino-acid sequences of *C. caldarium* PSI and LHCI; T.S. and N.D. verified polypeptide compositions; M.K. and K.I. performed phylogenetic analysis; T.H. collected cryo-EM images; Koji.K. processed the cryo-EM data and reconstructed the final cryo-EM map; Koji.K. built the structural model and refined the final model; Keisuke.K. contributed to the interpretation of the structure; Koji.K., Y.N., and R.N. evaluated the structural data; K.Y. and J.-R.S. provided experimental environments; R.N., Koji.K., T.H., M.K., K.I., S.H., S.-y.M., and K.Y. drafted the original manuscript; R.N. and J.-R.S. revised the manuscript; and R.N. wrote the final manuscript, and all of the authors joined the discussion of the results.

## References

1 Blankenship, R. E. Molecular Mechanisms of Photosynthesis. 3rd edn, (Wiley-Blackwell, 2021).

2 Green, B. R. & Durnford, D. G. The chlorophyll-carotenoid proteins of oxygenic photosynthesis. Annu. Rev. Plant Physiol. Plant Mol. Biol. 47, 685–714 (1996).

3 Shen, J.-R. in Macromolecular Protein Complexes IV. Subcellular Biochemistry (eds Harris, J. R. & Marles-Wright, J.) 351–377 (Springer, 2022).

4 Falkowski, P. G. et al. The evolution of modern eukaryotic phytoplankton. Science 305, 354–360 (2004).

5 Hippler, M. & Nelson, N. The plasticity of photosystem I. Plant Cell Physiol. 62, 1073–1081 (2021).

6 Antoshvili, M., Caspy, I., Hippler, M. & Nelson, N. Structure and function of photosystem I in *Cyanidioschyzon merolae*. Photosynth. Res. 139, 499–508 (2019).

7 Pi, X. et al. Unique organization of photosystem I-light-harvesting supercomplex revealed by cryo-EM from a red alga. Proc. Natl. Acad. Sci. U. S. A. 115, 4423–4428 (2018).

8 You, X. et al. In situ structure of the red algal phycobilisome-PSII-PSI-LHC megacomplex. Nature 616, 199–206 (2023).

9 Nagao, R. et al. Structural basis for assembly and function of a diatom photosystem I-light-harvesting supercomplex. Nat. Commun. 11, 2481 (2020).

10 Xu, C. et al. Structural basis for energy transfer in a huge diatom PSI-FCPI supercomplex. Nat. Commun. 11, 5081 (2020).

11 Zhao, L.-S. et al. Structural basis and evolution of the photosystem I-light-harvesting supercomplex of cryptophyte algae. Plant Cell (2023).

12 Yoon, H. S., Zuccarello, G. C. & Bhattacharya, D. in Red Algae in the Genomic Age (eds Seckbach, J. & Chapman, D. J.) 25–42 (Springer, 2010).

13 Ott, F. D. & Seckbach, J. in Evolutionary Pathways and Enigmatic Algae: Cyanidium caldarium (Rhodophyta) and Related Cells. (ed Seckbach, J.) 145–152 (Springer, 1994).

14 Ott, F. D. Handbook of the taxonomic names associated with the non-marine Rhodophycophyta. (J. Cramer, 2009).

15 Liu, S.-L., Chiang, Y.-R., Yoon, H. S. & Fu, H.-Y. Comparative genome analysis reveals *Cyanidiococcus* gen. nov., A new extremophilic red algal genus sister to *Cyanidioschyzon* (Cyanidioschyzonaceae, Rhodophyta). J. Phycol. 56, 1428–1442 (2020).

16 Park, S. I. et al. Revised classification of the Cyanidiophyceae based on plastid genome data with descriptions of the Cavernulicolales ord. nov. and Galdieriales ord. nov. (Rhodophyta). J. Phycol. 59, 444–466 (2023).

17 Tajima, N. et al. Analysis of the complete plastid genome of the unicellular red alga *Porphyridium purpureum*. J. Plant Res. 127, 389–397 (2014).

18 Engelken, J., Brinkmann, H. & Adamska, I. Taxonomic distribution and origins of the extended LHC (light-harvesting complex) antenna protein superfamily. BMC Evol. Biol. 10, 233 (2010).

19 Sturm, S. et al. A novel type of light-harvesting antenna protein of red algal origin in algae with secondary plastids. BMC Evol. Biol. 13, 159 (2013).

20 Ciniglia, C., Yoon, H. S., Pollio, A., Pinto, G. & Bhattacharya, D. Hidden biodiversity of the extremophilic Cyanidiales red algae. Mol. Ecol. 13, 1827–1838 (2004).

21 Gardian, Z. et al. Organisation of Photosystem I and Photosystem II in red alga *Cyanidium caldarium*: Encounter of cyanobacterial and higher plant concepts. Biochim. Biophys. Acta, Bioenerg. 1767, 725–731 (2007).

22 Nagao, R. et al. Biochemical and spectroscopic characterization of PSI-LHCI from the red alga *Cyanidium caldarium*. Photosynth. Res. 156, 315–323 (2023).

23 Nagao, R., Ueno, Y., Akimoto, S. & Shen, J.-R. Effects of CO_2_ and temperature on photosynthetic performance in the diatom *Chaetoceros gracilis*. Photosynth. Res. 146, 189–195 (2020).

24 Kumazawa, M. et al. Molecular phylogeny of fucoxanthin-chlorophyll *a*/*c* proteins from *Chaetoceros gracilis* and Lhcq/Lhcf diversity. Physiol. Plant. 174, e13598 (2022).

25 Kim, J. I. et al. Evolutionary dynamics of cryptophyte plastid genomes. Genome Biol. Evol. 9, 1859–1872 (2017).

26 Matsuzaki, M. et al. Genome sequence of the ultrasmall unicellular red alga *Cyanidioschyzon merolae* 10D. Nature 428, 653–657 (2004).

27 Cho, C. H. et al. Genome-wide signatures of adaptation to extreme environments in red algae. Nat. Commun. 14, 10 (2023).

28 Yonekura, K., Maki-Yonekura, S., Naitow, H., Hamaguchi, T. & Takaba, K. Machine learning-based real-time object locator/evaluator for cryo-EM data collection. Commun. Biol. 4, 1044 (2021).

29 Mastronarde, D. N. Automated electron microscope tomography using robust prediction of specimen movements. J. Struct. Biol. 152, 36–51 (2005).

30 Martin, M. Cutadapt removes adapter sequences from high-throughput sequencing reads. EMBnet J. 17, 10–12 (2011).

31 Bankevich, A. et al. SPAdes: a new genome assembly algorithm and its applications to single-cell sequencing. J. Comput. Biol. 19, 455–477 (2012).

32 Zheng, S. Q. et al. MotionCor2: anisotropic correction of beam-induced motion for improved cryo-electron microscopy. Nat. Methods 14, 331–332 (2017).

33 Mindell, J. A. & Grigorieff, N. Accurate determination of local defocus and specimen tilt in electron microscopy. J. Struct. Biol. 142, 334–347 (2003).

34 Zivanov, J., Nakane, T. & Scheres, S. H. W. Estimation of high-order aberrations and anisotropic magnification from cryo-EM data sets in *RELION*-3.1. IUCrJ 7, 253–267 (2020).

35 Grigorieff, N. & Harrison, S. C. Near-atomic resolution reconstructions of icosahedral viruses from electron cryo-microscopy. Curr. Opin. Struc. Biol. 21, 265–273 (2011).

36 Bepler, T., Kelley, K., Noble, A. J. & Berger, B. Topaz-Denoise: general deep denoising models for cryoEM and cryoET. Nat. Commun. 11, 5208 (2020).

37 Kelley, L. A., Mezulis, S., Yates, C. M., Wass, M. N. & Sternberg, M. J. E. The Phyre2 web portal for protein modeling, prediction and analysis. Nat. Protoc. 10, 845–858 (2015).

38 Pettersen, E. F. et al. UCSF Chimera - A visualization system for exploratory research and analysis. J. Comput. Chem. 25, 1605–1612 (2004).

39 Emsley, P., Lohkamp, B., Scott, W. G. & Cowtan, K. Features and development of *Coot*. Acta Crystallogr. D Biol. Crystallogr. 66, 486–501 (2010).

40 Adams, P. D. et al. PHENIX: a comprehensive Python-based system for macromolecular structure solution. Acta Crystallogr. D Biol. Crystallogr. 66, 213–221 (2010).

41 Yamashita, K., Palmer, C. M., Burnley, T. & Murshudov, G. N. Cryo-EM single-particle structure refinement and map calculation using Servalcat. Acta Crystallogr. D Struct. Biol. 77, 1282–1291 (2021).

42 Chen, V. B. et al. MolProbity: all-atom structure validation for macromolecular crystallography. Acta Crystallogr. D Biol. Crystallogr. 66, 12–21 (2010).

43 Barad, B. A. et al. EMRinger: side chain-directed model and map validation for 3D cryo-electron microscopy. Nat. Methods 12, 943–946 (2015).

44 Pintilie, G. et al. Measurement of atom resolvability in cryo-EM maps with *Q*-scores. Nat. Methods 17, 328–334 (2020).

45 Schrödinger, L. L. C. The PyMOL Molecular Graphics System. Version 2.5.0. (2021).

46 Pettersen, E. F. et al. UCSF ChimeraX: structure visualization for researchers, educators, and developers. Protein Sci. 30, 70–82 (2021).

47 Katoh, K. & Standley, D. M. MAFFT multiple sequence alignment software version 7: improvements in performance and usability. Mol. Biol. Evol. 30, 772–780 (2013).

48 Minh, B. Q. et al. IQ-TREE 2: new models and efficient methods for phylogenetic inference in the genomic era. Mol. Biol. Evol. 37, 1530–1534 (2020).

49 Kalyaanamoorthy, S., Minh, B. Q., Wong, T. K. F., von Haeseler, A. & Jermiin, L. S. ModelFinder: fast model selection for accurate phylogenetic estimates. Nat. Methods 14, 587–589 (2017).

50 Letunic, I. & Bork, P. Interactive Tree Of Life (iTOL) v5: an online tool for phylogenetic tree display and annotation. Nucleic Acids Res. 49, W293–W296 (2021).

51 Hoang, D. T., Chernomor, O., von Haeseler, A., Minh, B. Q. & Vinh, L. S. UFBoot2: improving the ultrafast bootstrap approximation. Mol. Biol. Evol. 35, 518–522 (2018).

