## Supplementary Information for "Structure of PSI-LHCI from *Cyanidium caldarium* provides evolutionary insights into conservation and diversity of red-lineage LHCs"

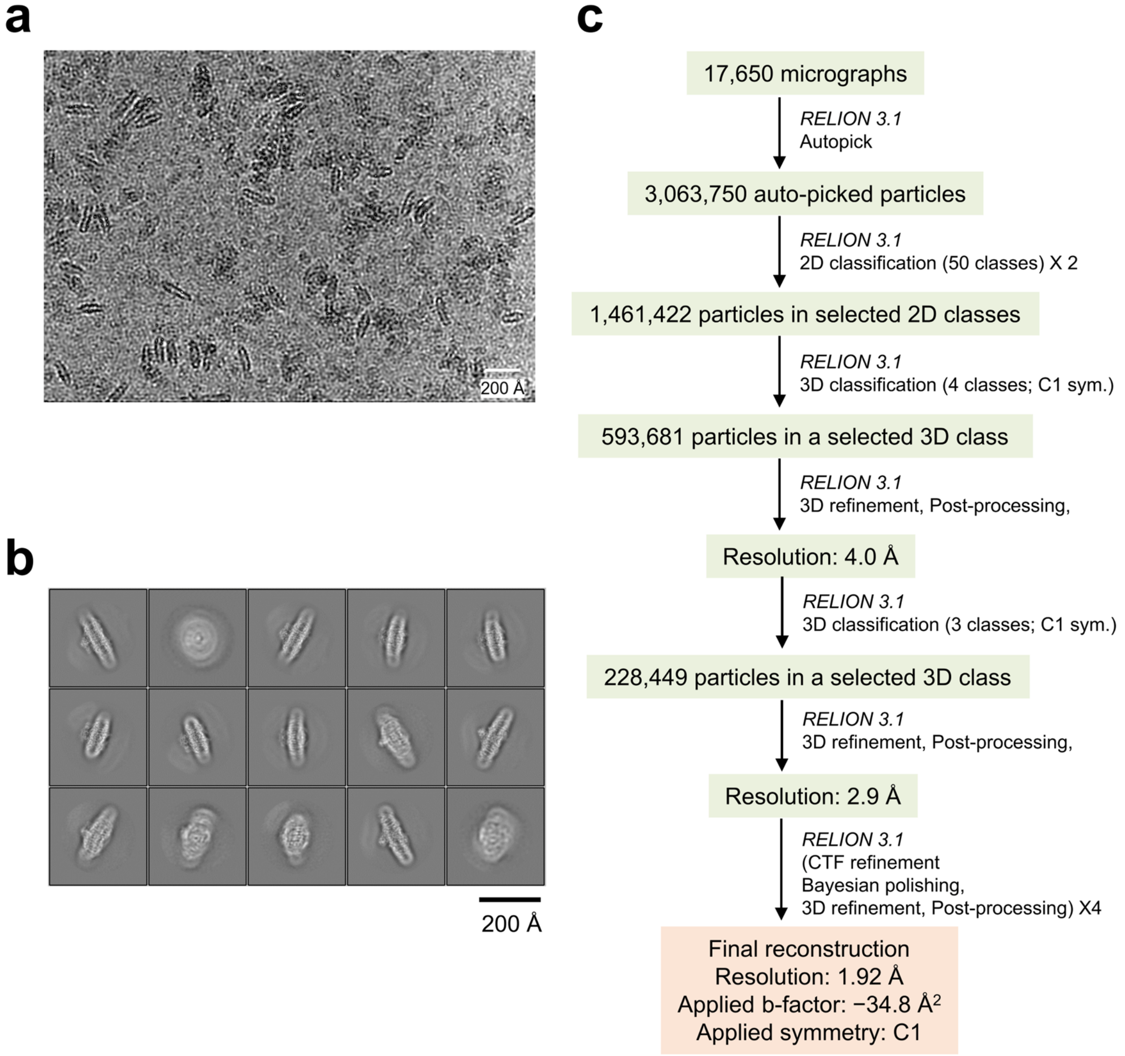


**Supplementary Fig. 1. Cryo-EM data collection and processing of PSI-LHCI.**

**a,** A representative cryo-EM micrograph of PSI-LHCI from 17,650 micrographs. **b,** Representative 2D classes of PSI-LHCI. The box size is 316.8 Å. **c,** A schematic flowchart showing the classification scheme and data processing for PSI-LHCI. The overall PSI-LHCI structure was reconstructed at a resolution of 1.92 Å from 228,449 particles. See Methods section for more details.

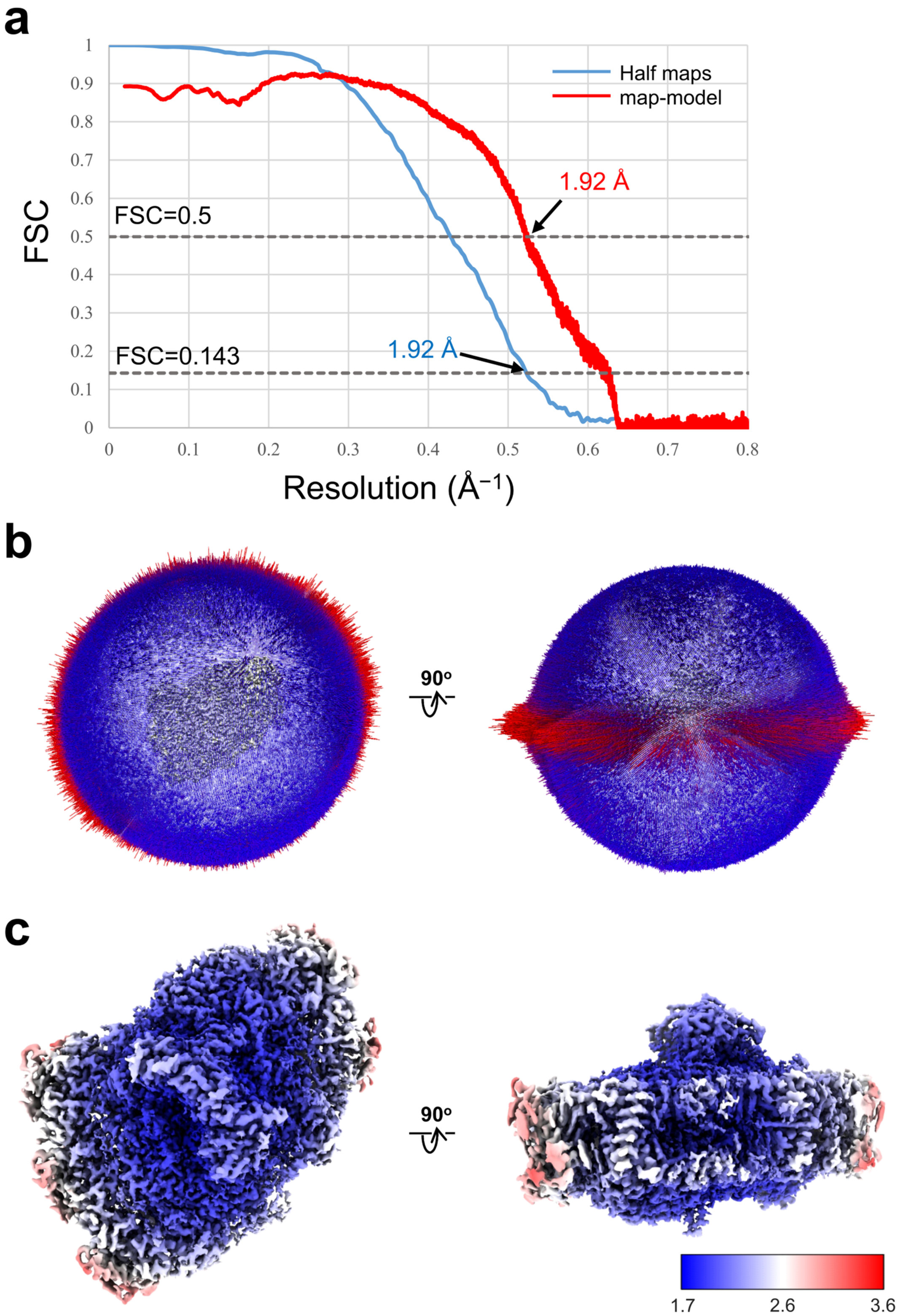


**Supplementary Fig. 2. Evaluation of the cryo-EM map quality.**

**a,** FSC curves of PSI-LHCI for independently refined half maps (blue) and map-minus-model (red). **b,** Angular distributions of the particles used for the reconstruction of PSI-LHCI. Each cylinder represents one view, and the height of the cylinder is proportional to the number of particles for that view. **c,** Local resolution maps of PSI-LHCI.

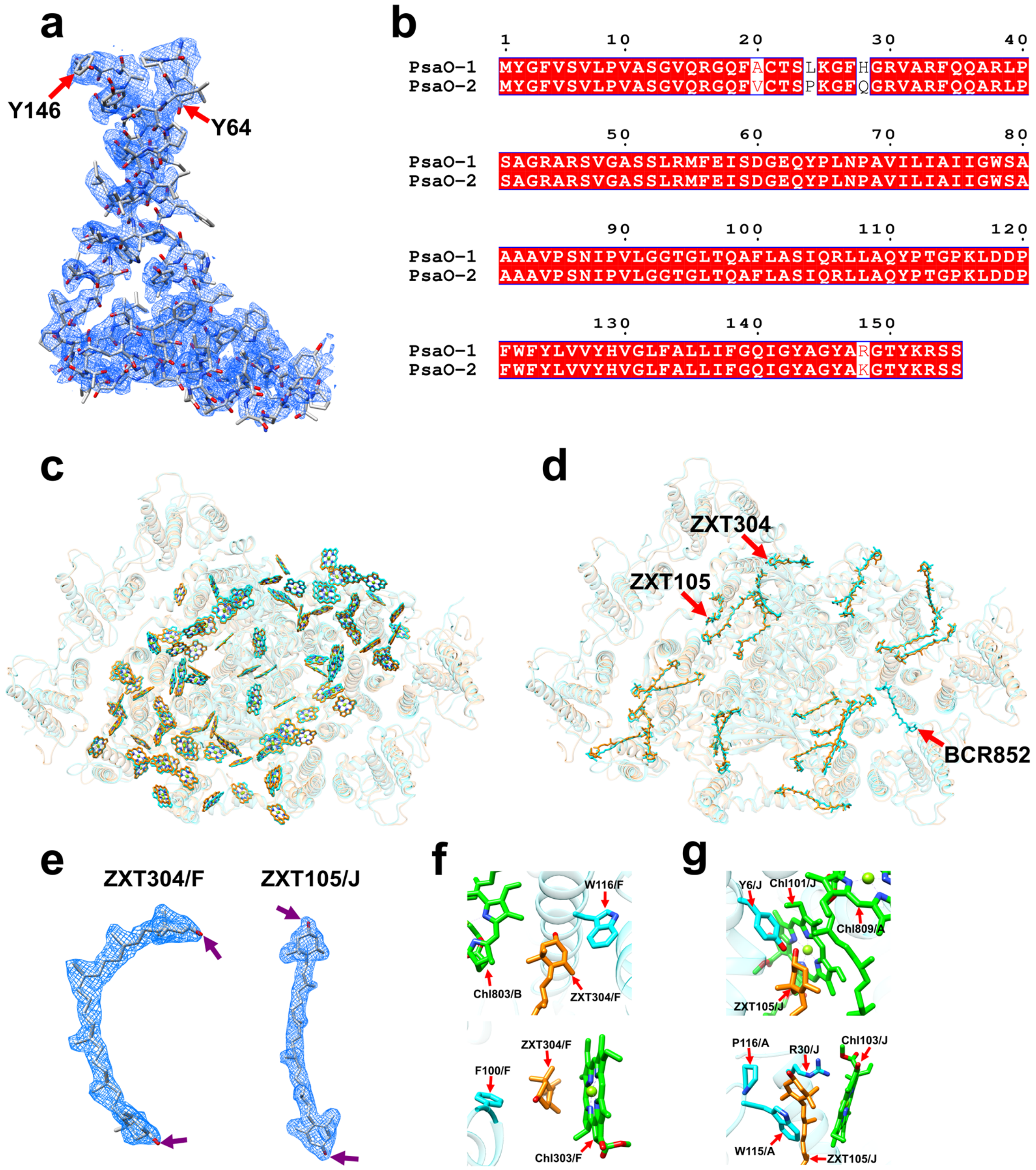


**Supplementary Fig. 3. Characteristic structure of the PSI core in the PSI-LHCI structure.** **a,** The cryo-EM density for PsaO and its corresponding model are shown as blue meshes and gray sticks, respectively. **b,** Multiple sequence alignment (ClustalW and ESPript) between PsaO-1 and PsaO-2 proteins. **c, d,** Comparison of Chls (**c**) and Cars (**d**) between *C. caldarium* (cyan) and *C. merolae* (orange) PSI-LHCIs. The *C. caldarium* PSI-LHCI structure overlapped with the *C. merolae* PSI-LHCI structure (PDB: 5ZGB). Chls and Cars are shown as sticks. Only rings of the Chl molecules are depicted. PsaB-BCR852 was found in the *C. caldarium* PSI-LHCI structure only, whereas PsaF-ZXT304 and PsaJ-ZXT105 in the *C. caldarium* PSI-LHCI correspond to BCR304 of PsaF and BCR105 of PsaJ, respectively, in the *C. merolae* PSI-LHCI. **e,** The densities and models of PsaF-ZXT304 and PsaJ-ZXT105 are shown as blue meshes and gray sticks, respectively. Purple arrows indicate the OH groups of ZXTs. **f, g,** Structures around the end rings of PsaF-ZXT304 (**f**) and PsaJ-ZXT105 (**g**). Proteins, Chls, and ZXTs are depicted in cyan, green, and orange, respectively.

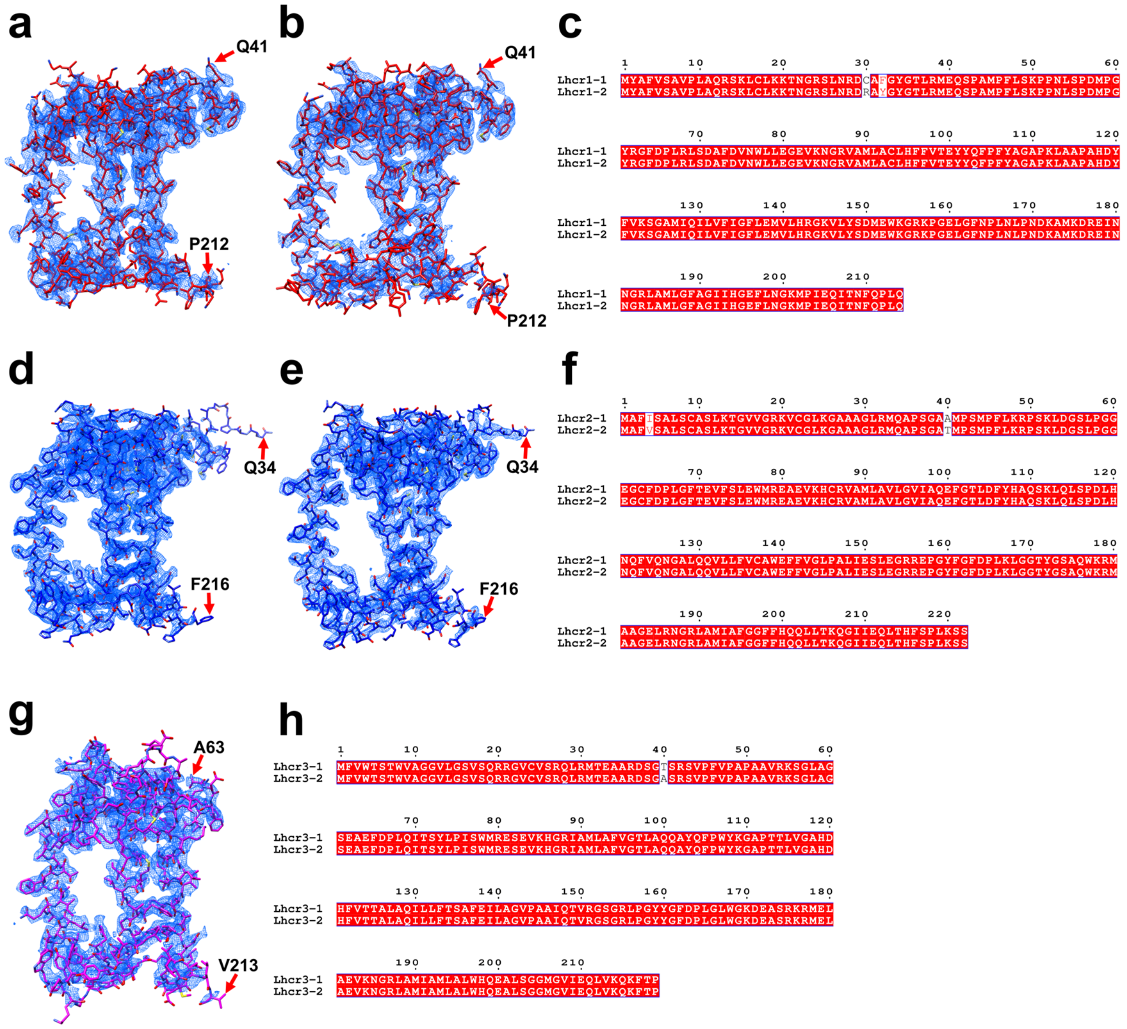


**Supplementary Fig. 4. Evaluation of the structures of LHCI subunits.**

**a, b,** The densities for LHCI-1 (**a**) and LHCI-3 (**b**), and their corresponding models are shown as meshes and sticks, respectively. **c,** Multiple sequence alignment (ClustalW and ESPript) between Lhcr1-1 and Lhcr1-2 proteins. **d, e,** The densities for LHCI-2 (**d**) and LHCI-4 (**e**), and their corresponding models are shown as meshes and sticks, respectively. **f,** Multiple sequence alignment (ClustalW and ESPript) between Lhcr2-1 and Lhcr2-2 proteins. **g,** The density for LHCI-3 and its corresponding model are shown as meshes and sticks, respectively. **h,** Multiple sequence alignment (ClustalW and ESPript) between Lhcr3-1 and Lhcr3-2 proteins.

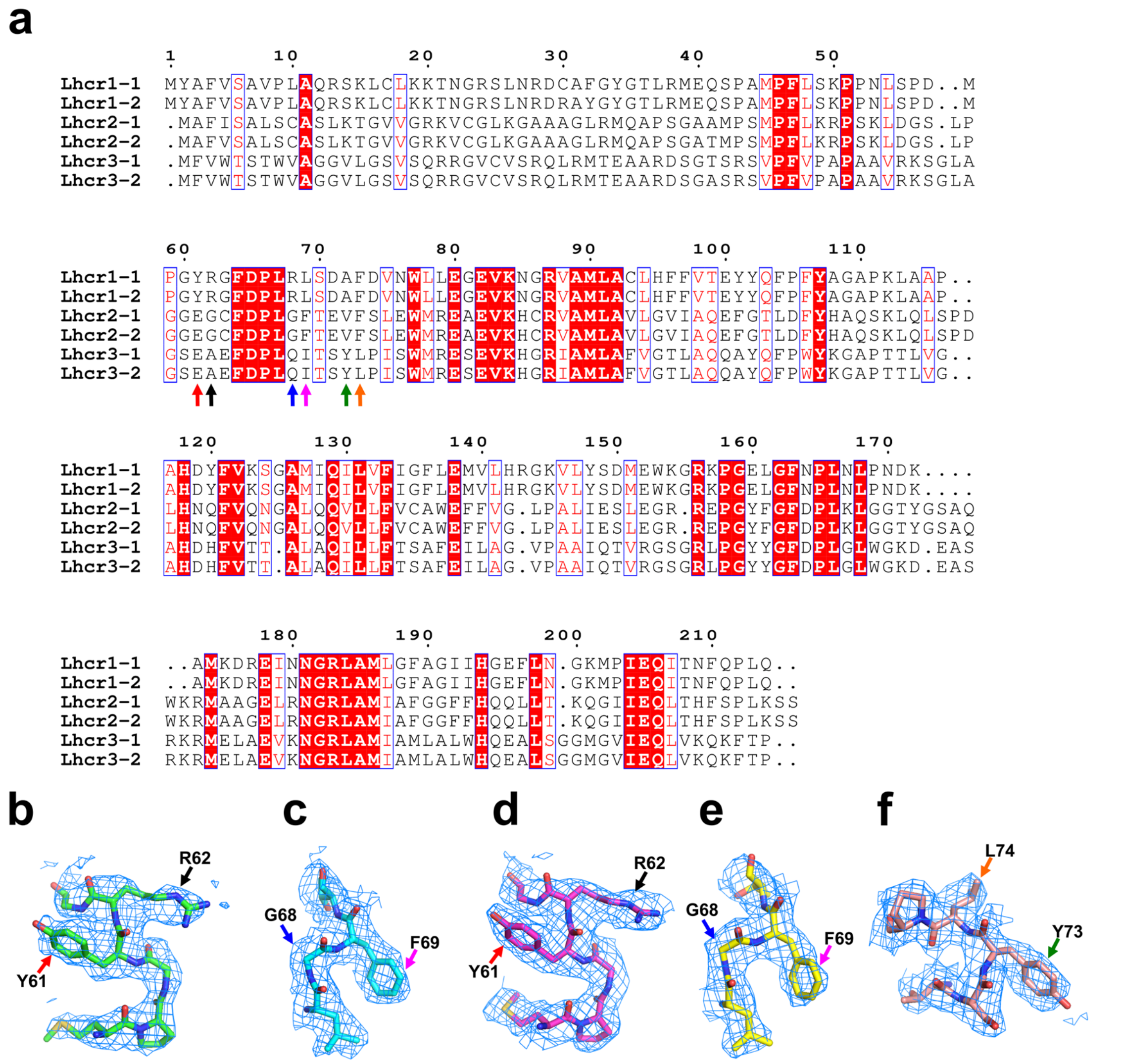


**Supplementary Fig. 5. Characteristic amino-acid residues used for the identification of different LHCI subunits.**

**a,** Multiple sequence alignment (ClustalW and ESPript) of Lhcrs of *C. caldarium*. Unique residues are indicated by arrows with different colors, which were used for the identification of different LHCIs. **b–f,** Characteristic maps and amino-acid residues of LHCI-1 (**b**), LHCI-2 (**c**), LHCI-3 (**d**), LHCI-4 (**e**), and LHCI-5 (**f**). The densities and models are shown as meshes and sticks, respectively. The characteristic amino acids are labeled with arrows of the same color shown in panel **a**.

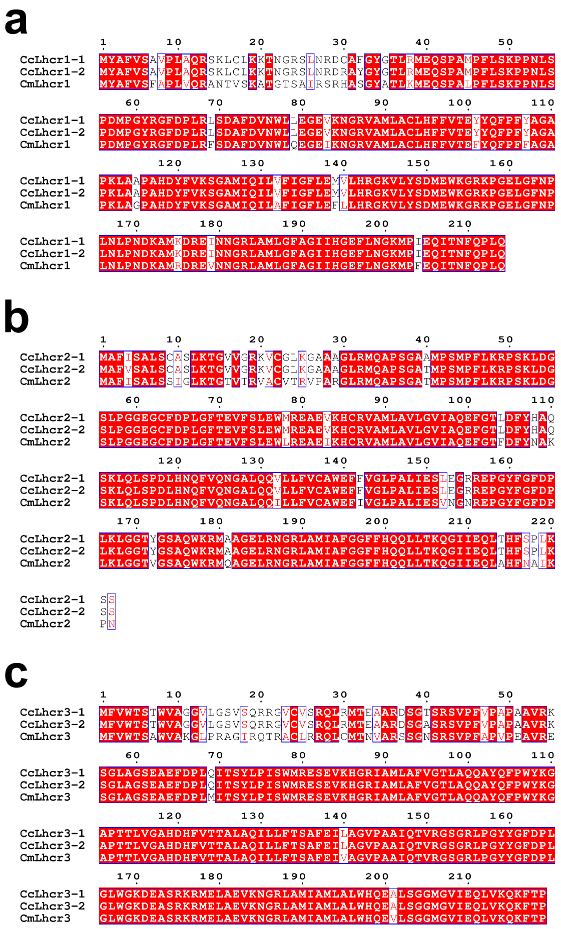

**Supplementary Fig. 6. Multiple sequence alignments of LHCI proteins between *C. caldarium* and *C. merolae*.**

Multiple sequence alignments of the LHC proteins located at the same positions in the PSI-LHCI structures of *C. caldarium* and *C. merolae* (PDB: 5ZGB) using ClustalW (https://www.genome.jp/tools-bin/clustalw) and ESPript (https://espript.ibcp.fr/ESPript/cgi-bin/ESPript.cgi). **a,** *C. caldarium* Lhcr1-1 (CcLhcr1-1), CcLhcr1-2, and *C. merolae* Lhcr1 (CmLhcr1). **b,** CcLhcr2-1, CcLhcr2-2, and CmLhcr2. **c,** CcLhcr3-1, CcLhcr3-2, and CmLhcr3. The gene IDs of CmLhcr1, CmLhcr2, and CmLhcr3 are CMQ142C, CMN234C, and CMN235C, respectively, in the genome database of *C. merolae* (http://czon.jp/). It should be noted that the sequence of CmLhcr3 was modified based on the *C. merolae* database and the PDB data of 5ZGB, because the open reading frame of CMN235C was interrupted in the genome.

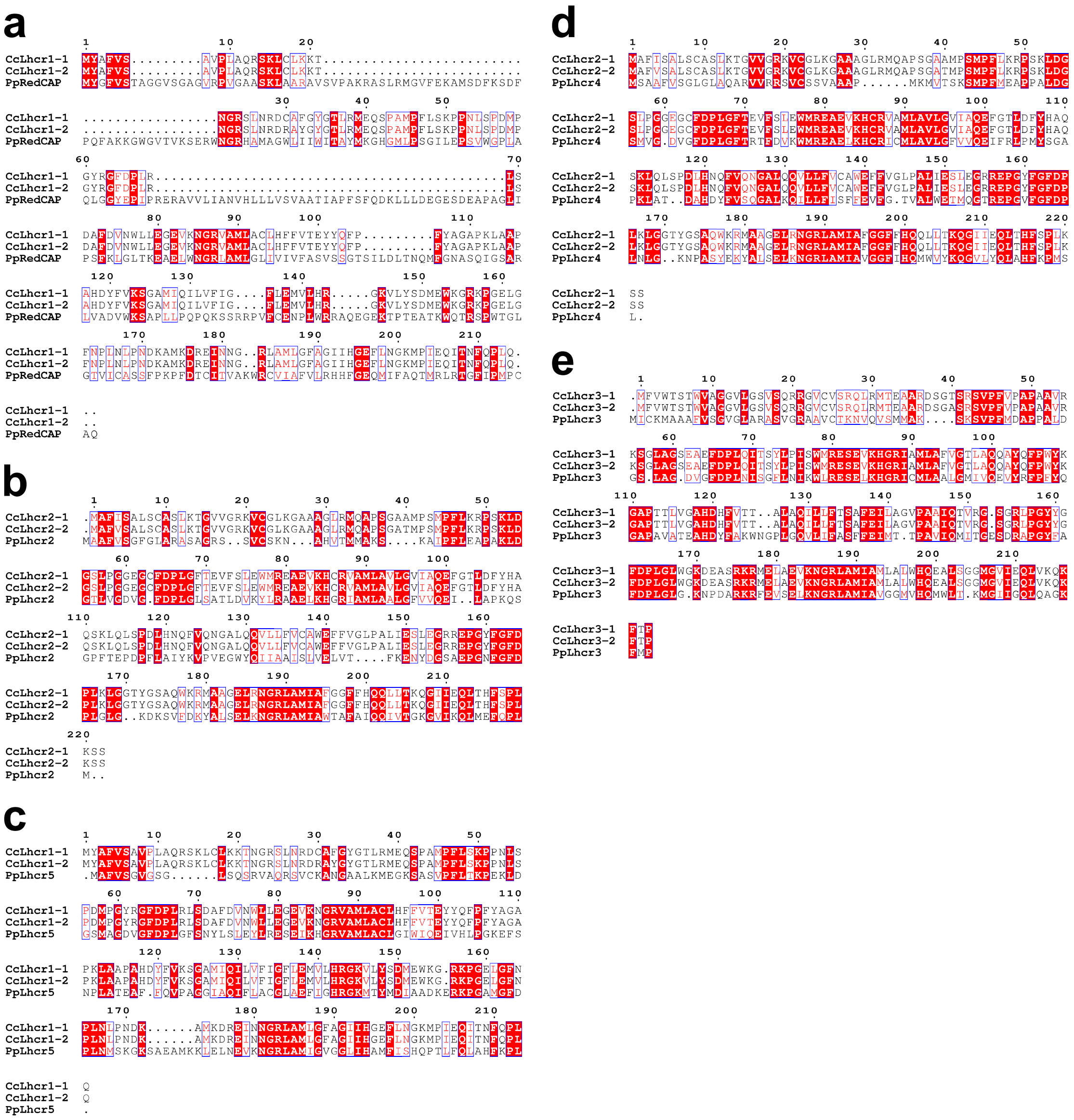

**Supplementary Fig. 7. Multiple sequence alignments of LHCI proteins between *C. caldarium* and *P. purpureum*.**

Multiple sequence alignments of the LHC proteins located at the same positions in the PSI-LHCI structures of *C. caldarium* and *P. purpureum* (PDB: 7Y5E) using ClustalW and ESPript. **a,** CcLhcr1-1, CcLhcr1-2, and *P. purpureum* RedCAP (PpRedCAP). **b,** CcLhcr2-1, CcLhcr2-2, and PpLhcr2. **c,** CcLhcr1-1, CcLhcr1-2, and PpLhcr5. **d,** CcLhcr2-1, CcLhcr2-2, and PpLhcr4. **e,** CcLhcr3-1, CcLhcr3-2, and PpLhcr3. The gene IDs of PpRedCAP, PpLhcr2, PpLhcr5, PpLhcr4, and PpLhcr3 are KAA8493301.1, KAA8498066.1, KAA8493980.1, KAA8499797.1, and KAA8496609.1, respectively, in the NCBI database.

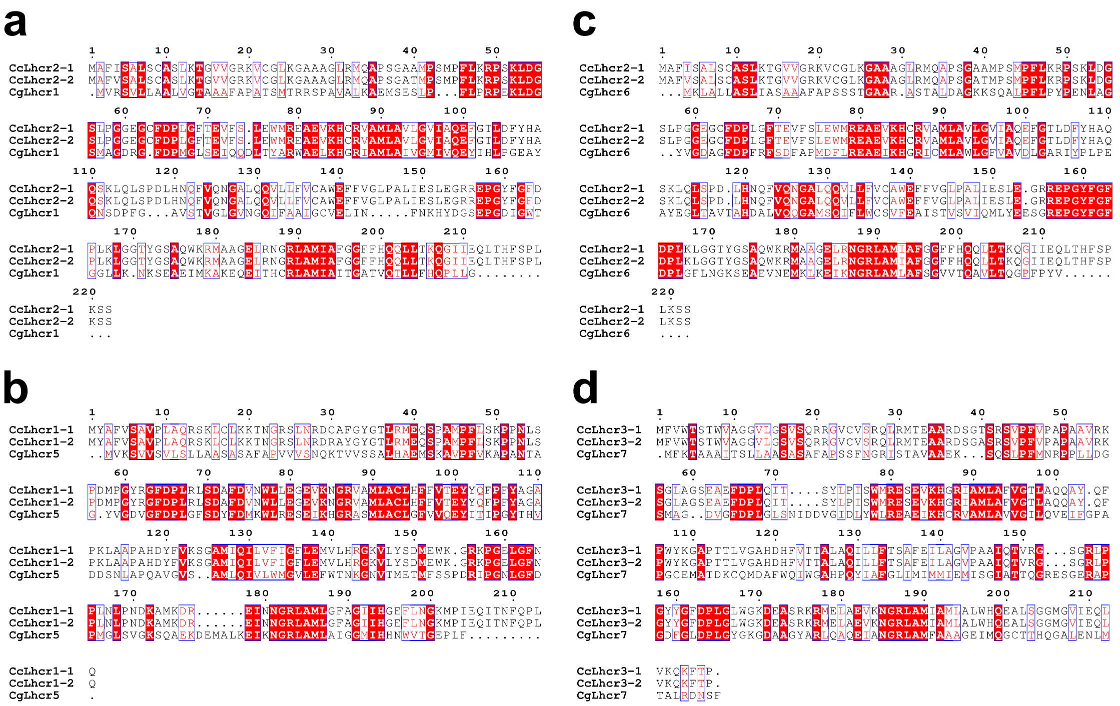

**Supplementary Fig. 8. Multiple sequence alignments of the *C. caldarium* LHCI proteins with the *C. gracilis* FCPIs.**

Multiple sequence alignments of the LHC proteins located at the same positions in the *C. caldarium* PSI-LHCI and *C. gracilis* PSI-FCPI (PDB: 6L4U) structures using ClustalW and ESPript. **a,** CcLhcr2-1, CcLhcr2-2, and *C. gracilis* Lhcr1 (CgLhcr1). **b,** CcLhcr1-1, CcLhcr1-2, and CgLhcr5. **c,** CcLhcr2-1, CcLhcr2-2, and CgLhcr6. **d,** CcLhcr3-1, CcLhcr3-2, and CgLhcr7. The gene IDs of CgLhcr1, CgLhcr5, CgLhcr6, and CgLhcr7 are g11656.t1, g11413.t1, g11475.t1, and g11027.t1, respectively, in the genome database of *C. gracilis* (https://chaetoceros.nibb.ac.jp/).

**a**

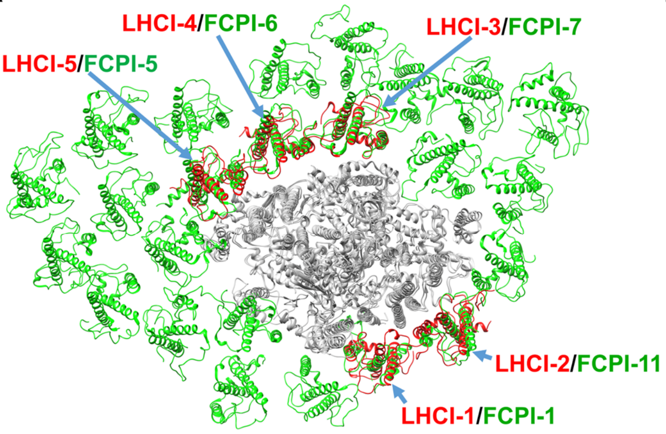


**b**

| *C. caldarium* | | *C. gracilis* | |
|---|---|---|---|
| Subunit | Gene | Subunit | Gene |
| LHCI-1 | *Lhcr1* | FCPI-1 | *fc13194* |
| LHCI-2 | *Lhcr2* | FCPI-11 | *lhcr3* |
| LHCI-3 | *Lhcr1* | FCPI-7 | *lhcr1* |
| LHCI-4 | *Lhcr2* | FCPI-6 | *lhcr4* |
| LHCI-5 | *Lhcr3* | FCPI-5 | *lhcr7* |

**c**

```
fc13194    ........................................................
CgRedCAP   MAPLRTLAALILGASAASAFAPTTPLATRNVVPAVSETSLNVFDKAMAEWEAEYPD

               1        10        20        30        40        50
fc13194    ....GWGPSVHAEKWNGRHAMFGWFFICCTAYAKGHGLIPDMDVPLNLKEWGTLAT
CgRedCAP   FAKWGWGPSVQAEKWNGRHAMFGWVFICATAYAKGHGLIPDADVPVDFSQWGTLAT

              60        70        80        90
fc13194    ITGKGTITNGRAVILLANAHFFAISLMATICPLPFGDSLLL...............
CgRedCAP   ISGKSTLTNERAIILIANVHCFLVGLMATICPLPFSDPLLLDPSQGEMYEKAKATK

                     100       110       120       130
fc13194    ...........LTEEAEMINGRLAMLGLISLIFATAIEQKPMLDIVNEWT.....
CgRedCAP   PYGIMPAMKMGLTEEAEIMNGRLAMMGIITVTATAFLQNKPILDVVNEWVGNAYY
```

**Supplementary Fig. 9. Structural comparison of the *C. caldarium* LHCIs with the *C. gracilis* FCPIs.**

**a,** Superposition of the *C. caldarium* PSI-LHCI structure with the *C. gracilis* PSI-FCPI structure with 24 FCPIs (PDB: 6LY5). The *C. caldarium* LHCI and *C. gracilis* FCPI subunits were colored red and green, respectively. The structures are viewed from the stromal side. **b,** Correlation of the names of LHCs in the structures with their genes between *C. caldarium* and *C. gracilis*. The names of LHCs and their genes are derived from Xu et al. (2020) for *C. gracilis*[10]. The gene of fc13194 is derived from a diatom *Fragilariopsis cylindrus*, whereas the genes of *lhcr3*, *lhcr1*, *lhcr4*, and *lhcr7* originate from transcriptome data of *C. gracilis* prepared by the authors of Xu et al. (2020)[10]. **c,** Sequence alignment of fc13124 with CgRedCAP (Gene ID: g6493.t1) using ClustalW and ESPript. The amino acid sequences of fc13124 and CgRedCAP were obtained from the PDB data of 6LY5 and the database of *C. gracilis*, respectively.

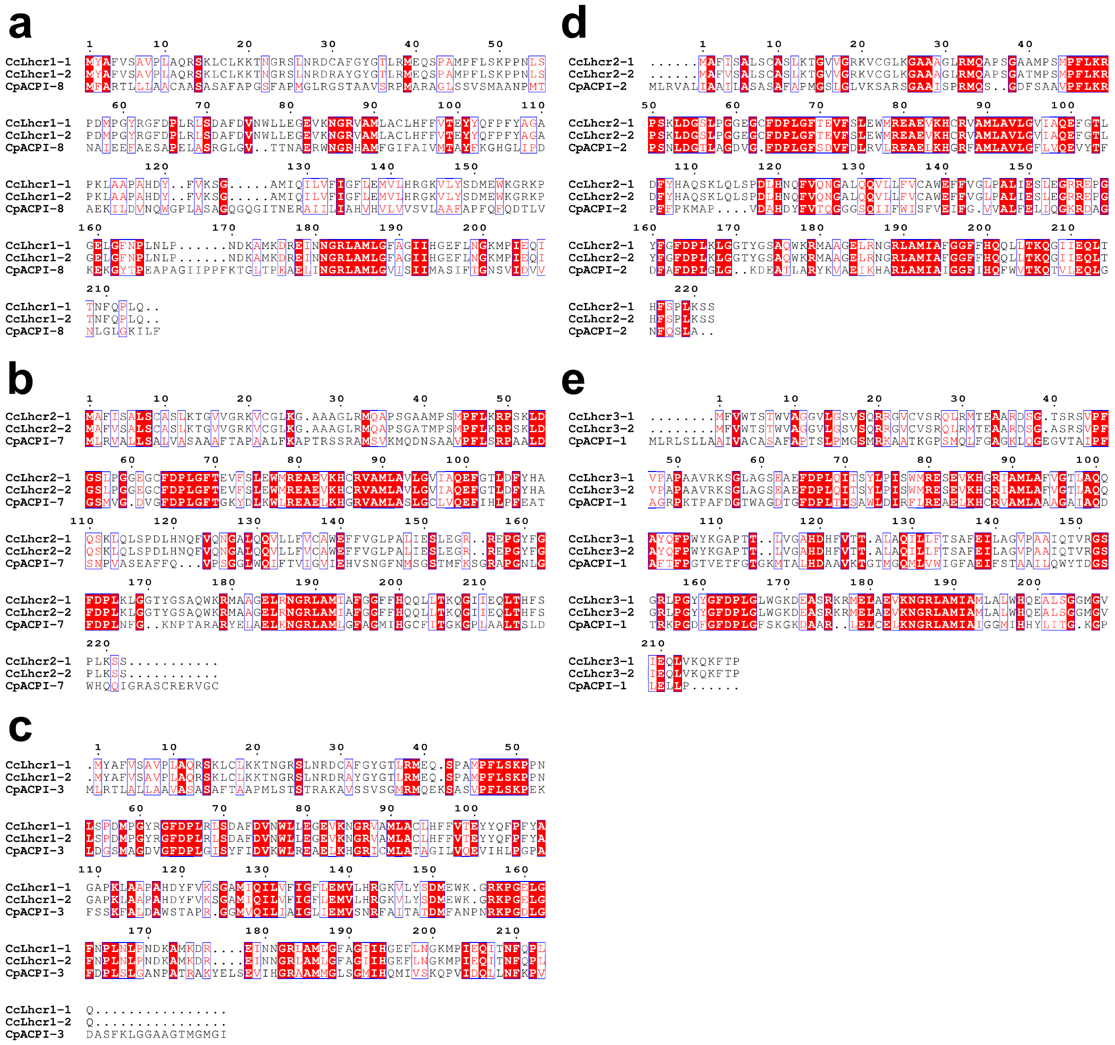

**Supplementary Fig. 10. Multiple sequence alignments of the *C. caldarium* LHCI proteins with the *C. placoidea* ACPIs.**

Multiple sequence alignments of the LHC proteins located at the same positions in the *C. caldarium* PSI-LHCI and *C. placoidea* PSI-ACPI (PDB: 7Y7B) structures using ClustalW and ESPript. **a,** CcLhcr1-1, CcLhcr1-2, and *C. placoidea* ACPI-8 (CpACPI-8). **b,** CcLhcr2-1, CcLhcr2-2, and CpACPI-7. **c,** CcLhcr1-1, CcLhcr1-2, and CpACPI-3. **d,** CcLhcr2-1, CcLhcr2-2, and CpACPI-2. **e,** CcLhcr3-1, CcLhcr3-2, and CpACPI-1. The ACPI sequences were based on the transcriptome data of *C. placoidea* by Zhao et al. (2023)[11].

**a**

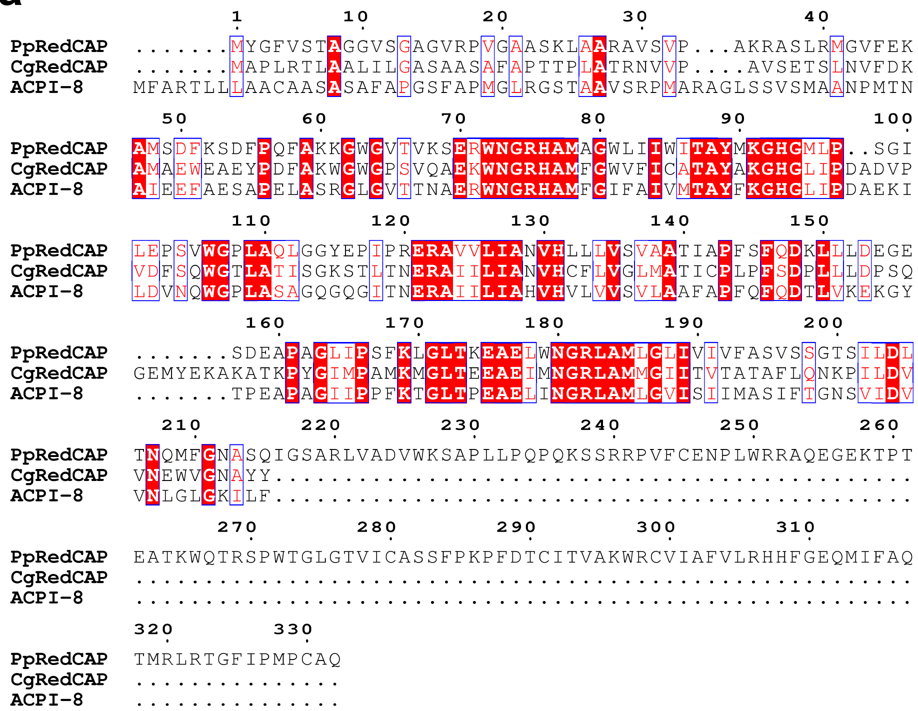


**Supplementary Fig. 11. Multiple sequence alignments of the *C. placoidea* ACPI-8 with the *P. purpureum* and *C. gracilis* RedCAPs.**

Multiple sequence alignments using ClustalW and ESPript. The amino acid sequences of PpRedCAP and CgRedCAP are derived from the gene products of KAA8493301.1 in *P. purpureum* and g6493.t1 in *C. gracilis*, respectively.

**Supplementary Table 1. Cryo-EM data collection and structural analysis statistics.**

| | |
|---|---|
| Complex | PSI-LHCI |
| PDB ID | 8WEY |
| EMDB ID | EMD-37480 |
| Data collection and processing | |
| Magnification | 100,000 |
| Voltage (kV) | 300 |
| Electron exposure (e–/Å) | 39.8 |
| Defocus range (μm) | −1.8 to −0.8 |
| Physical pixel size (Å) | 0.495 |
| Symmetry imposed | C1 |
| Initial particle images (no.) | 3,063,750 |
| Final particle images (no.) | 228,449 |
| Map resolution (Å) | 1.92 |
| FSC threshold | 0.143 |
| Refinement | |
| Initial model used | Homology modeling |
| Model resolution (Å) | 2.14 |
| FSC threshold | 0.5 |
| Map sharpening B factor ($Å^2$) | −34.8 |
| Model composition | |
| Non-hydrogen atoms | 36,163 |
| Protein | 24,460 |
| Ligand | 11,143 |
| Water | 560 |
| B factors ($Å^2$) | |
| Protein | 67.4 |
| Ligand | 76.0 |
| Water | 44.2 |
| R.m.s deviations | |
| Bond lengths (Å) | 0.028 |
| Bond angles (°) | 2.89 |
| Validation | |
| MolProbity score | 1.83 |
| Clashscore | 7.33 |
| Poor rotamers (%) | 1.34 |
| EMRinger score | 5.85 |
| Ramachandran plot | |
| Favored (%) | 95.26 |
| Allowed (%) | 4.32 |
| Disallowed (%) | 0.42 |

**Supplementary Table 2. Averaged *Q*-score in each subunit.**

| Subunit | Averaged *Q*-score | |
|---|---|---|
| | Postprocessed map | Denoised map |
| PsaA | 0.87 | 0.88 |
| PsaB | 0.86 | 0.87 |
| PsaC | 0.90 | 0.90 |
| PsaD | 0.81 | 0.83 |
| PsaE | 0.81 | 0.84 |
| PsaF | 0.82 | 0.84 |
| PsaI | 0.82 | 0.84 |
| PsaJ | 0.86 | 0.87 |
| PsaK | 0.69 | 0.75 |
| PsaL | 0.72 | 0.76 |
| PsaM | 0.80 | 0.82 |
| PsaO | 0.38 | 0.56 |
| LHCI-1 | 0.44 | 0.59 |
| LHCI-2 | 0.62 | 0.70 |
| LHCI-3 | 0.54 | 0.64 |
| LHCI-4 | 0.58 | 0.69 |
| LHCI-5 | 0.53 | 0.66 |

**Supplementary Table 3. Cofactors assigned in the PSI-LHCI structure.**

| Protein | Chlorophyll | Carotenoid | Lipid | Other |
|---|---|---|---|---|
| PsaA | 39 Chl *a*<br>1 Chl *a′* | 4 BCR | 2 LHG | 1 [4Fe-4S] cluster,<br>1 phylloquinone |
| PsaB | 42 Chl *a* | 8 BCR | 1 LHG<br>1 DGD | 1 phylloquinone |
| PsaC | - | - | - | 2 [4Fe-4S] cluster |
| PsaD | - | - | - | - |
| PsaE | - | - | - | - |
| PsaF | 3 Chl *a* | 1 ZXT | - | - |
| PsaI | - | 1 BCR | - | - |
| PsaJ | 3 Chl *a* | 1 BCR<br>1 ZXT | - | - |
| PsaK | 2 Chl *a* | 2 BCR | - | - |
| PsaL | 3 Chl *a* | 3 BCR | - | - |
| PsaM | - | - | - | - |
| PsaO | 4 Chl *a* | 1 BCR | - | - |
| LHCI-1 | 9 Chl *a* | 4 ZXT | - | - |
| LHCI-2 | 13 Chl *a* | 4 ZXT | - | - |
| LHCI-3 | 11 Chl *a* | 5 ZXT | - | - |
| LHCI-4 | 13 Chl *a* | 4 ZXT | - | - |
| LHCI-5 | 11 Chl *a* | 4 ZXT | - | - |
| Total | 154 | 43 | 4 | 5 |

BCR, *β*-carotene; ZXT, zeaxanthin; Chl *a*, chlorophyll *a*; Chl *a′*, chlorophyll *a* epimer; DGD, digalactosyldiacyl glycerol; LHG, dipalmitoylphosphatidyl glycerol.

**Supplementary Table 4. Chls and their ligands in LHCIs.**

| Protein | Chlorophyll/ligand |
|---|---|
| LHCI-1 | a301/A44, a302/E82, a303/N85, a306/H118, a307/E138, a308/E178, a309/-[1], a310/N181, a311/-[1] |
| LHCI-2 | a301/S43, a302/E82, a303/H85, a304/Q99, a305/Q131, a306/H120, a307/E140, a308/E184, a309/-[1], a310/N187, a311/Q201, a312/H200, a313/-[1] |
| LHCI-3 | a301/A44, a302/E82, a303/N85, a304/H95, a306/H118, a307/E138, a308/E178, a309/-[1], a310/N181, a311/-[1], a312/H194 |
| LHCI-4 | a301/S43, a302/E82, a303/H85, a304/Q99, a305/Q131, a306/H120, a307/E140, a308/E184, a309/-[1], a310/N187, a311/Q201, a312/H200, a313/-[1] |
| LHCI-5 | a303/E83, a304/H86, a305/-[1], a306/Q129, a307/H119, a308/E138, a309/E182, a310/-[1], a311/N185, a312/Q199, a313/H198 |

[1]The ligands of these Chls may be water molecules which could not be identified due to weak densities.

**Supplementary Table 5. LHCI proteins identified in the *C. caldarium* PSI-LHCI structure and their RMSD values with LHCI-3**

| Protein | Gene* | RMSD (Å)/Aligned Cα atoms |
|---|---|---|
| LHCI-1 | *Lhcr1-1*/*Lhcr1-2* | 0.74/170 |
| LHCI-2 | *Lhcr2-1*/*Lhcr2-2* | 1.53/154 |
| LHCI-3 | *Lhcr1-1*/*Lhcr1-2* | 0.00/172 |
| LHCI-4 | *Lhcr2-1*/*Lhcr2-2* | 1.55/154 |
| LHCI-5 | *Lhcr3-1*/*Lhcr3-2* | 1.49/126 |

*These genes cannot be distinguished in the present structure.

**Supplementary Table 6. Correspondence of numbering of pigments in each PSI subunit described in the text with those in the PDB file.**

| | PsaF | PsaJ |
|---|---|---|
| **Chls in the text** | **PDB No. (Chain ID)** | **PDB No. (Chain ID)** |
| 101 | | 856 (A)* |
| 102 | | 857 (B)* |
| 103 | | 103 (J) |
| 301 | 855 (A)* | |
| 302 | 203 (F) | |
| 303 | 204 (F) | |
| **Cars in the text** | | |
| 104 | | 104 (J) |
| 105 | | 105 (J) |
| 304 | 205 (F) | |

*Chain in the adjacent unit.

**Supplementary Table 7. Correspondence of numbering of pigments in each LHCI subunit described in the text with those in the PDB file.**

| | LHCI-1 | LHCI-2 | LHCI-3 | LHCI-4 | LHCI-5 |
|---|---|---|---|---|---|
| **Chls in the text** | **PDB No. (Chain ID)** | **PDB No. (Chain ID)** | **PDB No. (Chain ID)** | **PDB No. (Chain ID)** | **PDB No. (Chain ID)** |
| 301 | 303 (4) | 302 (5) | 301 (1) | 304 (2) | |
| 302 | 304 (4) | 303 (5) | 302 (1) | 305 (2) | |
| 303 | 305 (4) | 304 (5) | 303 (1) | 306 (2) | 305 (3) |
| 304 | | 305 (5) | 304 (1) | 307 (2) | 306 (3) |
| 305 | | 306 (5) | | 308 (2) | 307 (3) |
| 306 | 306 (4) | 307 (5) | 305 (1) | 309 (2) | 308 (3) |
| 307 | 307 (4) | 308 (5) | 306 (1) | 310 (2) | 309 (3) |
| 308 | 308 (4) | 309 (5) | 307 (1) | 311 (2) | 310 (3) |
| 309 | 309 (4) | 310 (5) | 308 (1) | 312 (2) | 311 (3) |
| 310 | 310 (4) | 311 (5) | 309 (1) | 313 (2) | 312 (3) |
| 311 | 311 (4) | 312 (5) | 310 (1) | 314 (2) | 313 (3) |
| 312 | | 313 (5) | 311 (1) | 315 (2) | 314 (3) |
| 313 | | 314 (5) | | 316 (2) | 315 (3) |
| **Cars in the text** | | | | | |
| 301 | | | | | 321 (2)* |
| 312 | 312 (4) | | | | |
| 313 | 313 (4) | | 312 (1) | | |
| 314 | 314 (4) | 315 (5) | 313 (1) | 317 (2) | 316 (3) |
| 315 | | 316 (5) | 314 (1) | 318 (2) | 317 (3) |
| 316 | | 317 (5) | 315 (1) | 319 (2) | |
| 317 | 301 (5)* | 318 (5) | 316 (1) | 320 (2) | 318 (3) |

*Chain in the adjacent unit.
